# Metalorian: *De Novo* Generation of Heavy Metal-Binding Peptides with Classifier-Guided Diffusion Sampling

**DOI:** 10.1101/2025.07.10.664242

**Authors:** Yinuo Zhang, Divya Srijay, Zachary Quinn, Pranam Chatterjee

## Abstract

Heavy metal contamination is a severe and ongoing global problem that demands selective and efficient chelation strategies for environmental remediation. As a solution, we introduce **Metalorian**, a conditional diffusion model that generates *de novo* heavy metal-binding peptides, guided by **MetaLATTE**, a multi-label classifier fine-tuned to include underrepresented metal classes. Leveraging both continuous protein embeddings and discrete metal-binding constraints, Metalorian produces peptides with controllable lengths and user-defined binding specificity while preserving essential physicochemical properties. Crucially, our data augmentation strategy enables robust classification and generation even with sparse training data for certain metals. We demonstrate the effectiveness of our approach through peptides designed for copper (Cu), cadmium (Cd), and zinc (Zn), which retain key features such as net charge and hydrophobicity, while significantly reducing sequence length and molecular weight compared to known metal-binding proteins. Metalorian-generated peptides exhibit stable conformations and favorable binding energetics, as confirmed through molecular dynamics simulations. To validate binding *in vitro*, we developed a streamlined SUMO-fusion expression and cleavage system coupled with ELISA-based quantification, confirming robust Cu and Zn binding by multiple Metalorian-generated peptides. Overall, our work establishes a foundational platform for engineering heavy metal-binding peptides tailored to diverse and underrepresented targets, and highlights the potential of well-trained continuous latent spaces for diffusion-based *de novo* peptide design.

## Introduction

Metals play a crucial role in natural systems, serving as co-factors for the functionality of enzymes and catalysts for chemical reactions and pharmaceutical production [Kostenkova et al., 2022]. Transition metals are particularly essential due to their wide range of oxidation states and involvement in various biological processes. Most of the proteins in the Protein Data Bank (PDB) are metalloproteins [Permyakov, 2021], underscoring the ubiquity of metal-protein interactions in biology. However, this fundamental dependence on metals creates vulnerability when toxic heavy metals such as cadmium (Cd), lead (Pb), and mercury (Hg) are present at high concentrations [Tchounwou et al., 2012, Chen et al., 2023b], affecting nearly 14 billion people worldwide [Hou et al., 2025]. These toxic non-essential heavy metals disrupt cellular functions by interfering with essential metal-dependent processes, often exhibiting higher affinity for sulfhydryl groups in functional metalloenzymes and sometimes possess structurally-similar functional groups compared with essential metals like zinc (Zn), copper (Cu), and iron (Fe)[Tripathi and Poluri, 2021]. Given that these toxic metals persist in ecosystems, there is a critical demand for effective chelating approaches to remediate heavy metal contamination [Dixit et al., 2015, Zheng et al., 2021]. While certain natural metalloproteins, such as metallothioneins, can bind a range of metals [Li et al., 2023], they are challenging to repurpose for broader applications due to constraints in size, folding, and specificity requirements.

This challenge necessitates the development of computational and experimental methods that enable rapid generation and testing of metal-binding peptides. Existing metal predictor tools, such as MIB2 [Lu et al., 2022], require structural information to predict binding sites, limiting their utility for disordered or uncharacterized metalloproteins [Abramson et al., 2024, Permyakov, 2021]. Meanwhile, current *de novo* metal-binding peptide design pipelines are limited, with previous attempts relying on initial scaffold selection and directed evolution, which significantly constrain the available search space [Learte-Aymamí et al., 2024, Zhang et al., 2021]. These shortcomings collectively highlight the need for computational approaches that can generate objective-oriented metal-binding peptides with controllable length, high specificity, and strong binding affinity for targeted metal categories, coupled with efficient experimental validation methods.

In this work, we introduce **Metalorian**, a co-evolving conditional diffusion model that integrates continuous protein embeddings and discrete sequence properties to design metal-binding peptides *de novo* [Lee et al., 2023, Schroff et al., 2015]. At the core of our approach is **MetaLATTE** (**M**etal binding predictor using protein **LA**nguage model la**T**en**T E**mbeddings), a multi-label classifier that guides Metalorian’s generative process toward sequences enriched in key metal-binding motifs. For experimental validation, we developed a streamlined SUMO-fusion expression and cleavage system, enabling direct ELISA-based quantification of peptide–metal interactions without requiring extensive purification or biopanning. Unlike traditional metalbinding characterization methods that often rely on laborious structural modeling or iterative selection [Yang et al., 2015, Zheng et al., 2021, Matys et al., 2020], our platform allows rapid, quantitative assessment of binding. We demonstrate that MetaLATTE accurately predicts metal-binding properties from sequence alone, while Metalorian generates diverse and valid binders, which we rigorously validate via molecular dynamics (MD) simulations and *in vitro* assays. Taken together, our integrated pipeline serves as a versatile, end-to-end platform for discovering novel heavy metal-binding peptides, offering potential solutions from bioremediation to industrial catalysis.

## Results

### MetaLATTE enables accurate classification of peptide binders for cognate heavy metals

A key prerequisite for Metalorian’s guided peptide generation is a reliable predictor that can evaluate metalbinding properties directly from sequence. However, existing metal-binding prediction tools like MIB2 rely on structural information [Lu et al., 2022, Aptekmann et al., 2022], limiting their use on disordered proteins, novel peptides, or sequences without resolved 3D structures. Moreover, they often fail to capture subtle, residue-level differences, such as single-site mutations, that drastically alter metal-binding behavior. To overcome these limitations, we developed MetaLATTE, a multi-label classifier trained on protein language model embeddings from ESM-2 [Lin et al., 2023], specifically designed to capture both global context and local binding motifs across 14 transition metals.

MetaLATTE’s architecture (Figure 1C) incorporates an attention-pooling layer and rotary position embeddings atop ESM-2’s representations, and is trained with a two-stage approach. The first stage uses class-balanced focal and F1 losses to learn general metal-binding tendencies across the dataset. Building on this foundation, the second stage introduces triplet loss over synthetically mutated sequences (such as alanine swaps at binding residues; see Figure 1B), enabling the model to distinguish fine-grained functional differences in binding that are critical for accurate predictions.

**Figure 1:**
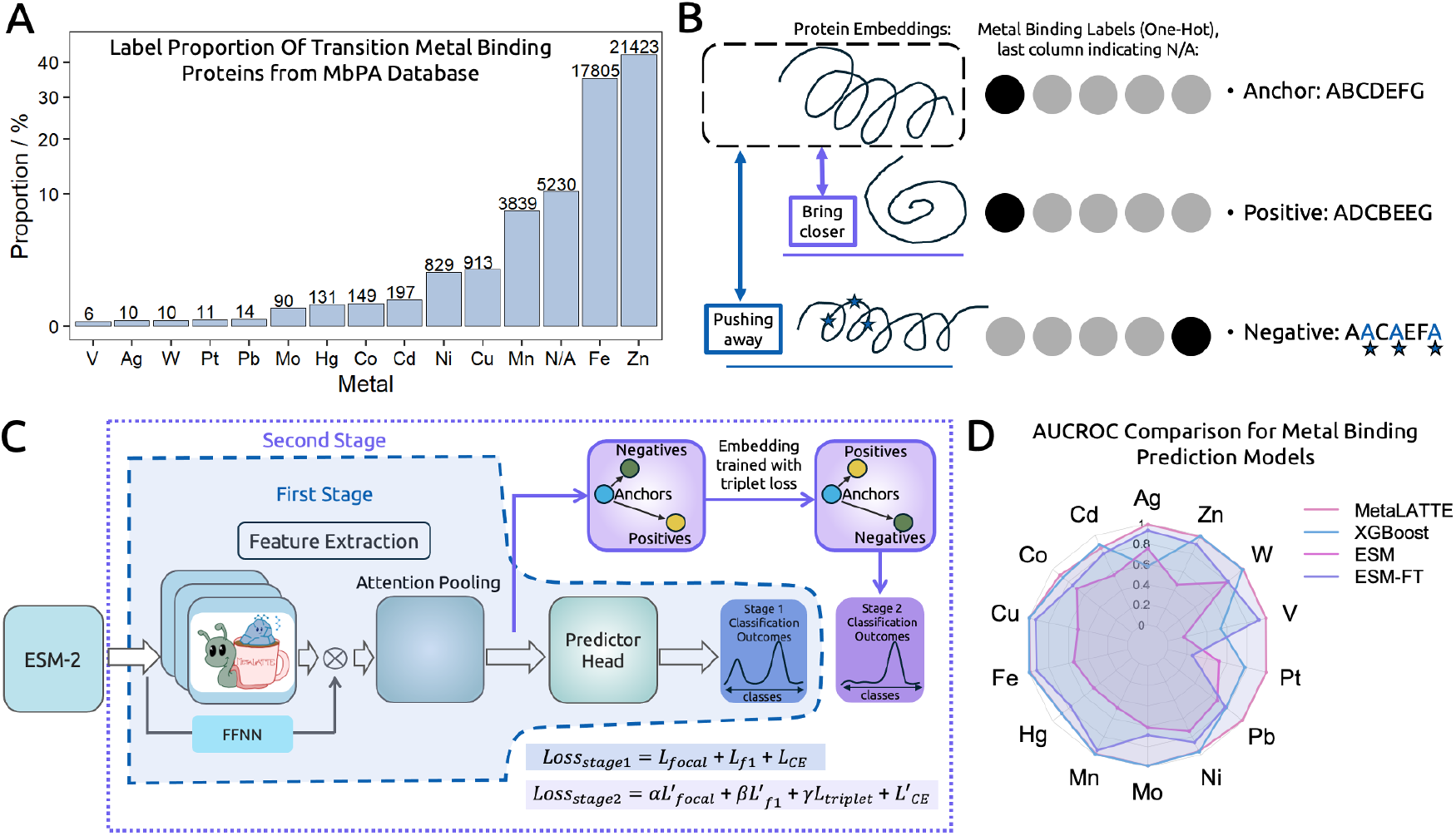
MetaLATTE schematic. **A)** Distribution of the metal-binding labels for the training data from MbPA database. **B)** Triplet loss is applied to the constructed pairs to differentiate the latent differences created by data augmentation. **C)** MetaLATTE schematic. Two stages of training is applied to fine-tune the ESM-2 model, and different loss combination is used during each stage. **D)** AUCROC performance over fourteen metal classes we included in this manuscript from our final model vs. the benchmark models. Our model, MetaLATTE, in generally performed well for all classes.

This training strategy proved highly effective, as MetaLATTE achieved high AUCROC scores (0.86-0.99) across all 14 metals, outperforming benchmark models particularly for rare metals like vanadium and silver (Figure 1D). The model demonstrated balanced performance with superior recall (0.55) and F1 score (0.57) compared to alternatives (Figure 2B). To understand how the two-stage training contributes to this performance, we analyzed the learned representations through UMAP visualization of triplet datasets, which revealed clear separation between anchor/positive proteins and their negative mutant clusters (Figure 2A). Such separation demonstrates MetaLATTE’s ability to detect binding changes due to mutations, confirming that while stage one focal loss alone can differentiate embeddings for different labels, the triplet training with introduced mutations around motif sites enables MetaLATTE to better distinguish functional changes due to amino acid mutations.

**Figure 2:**
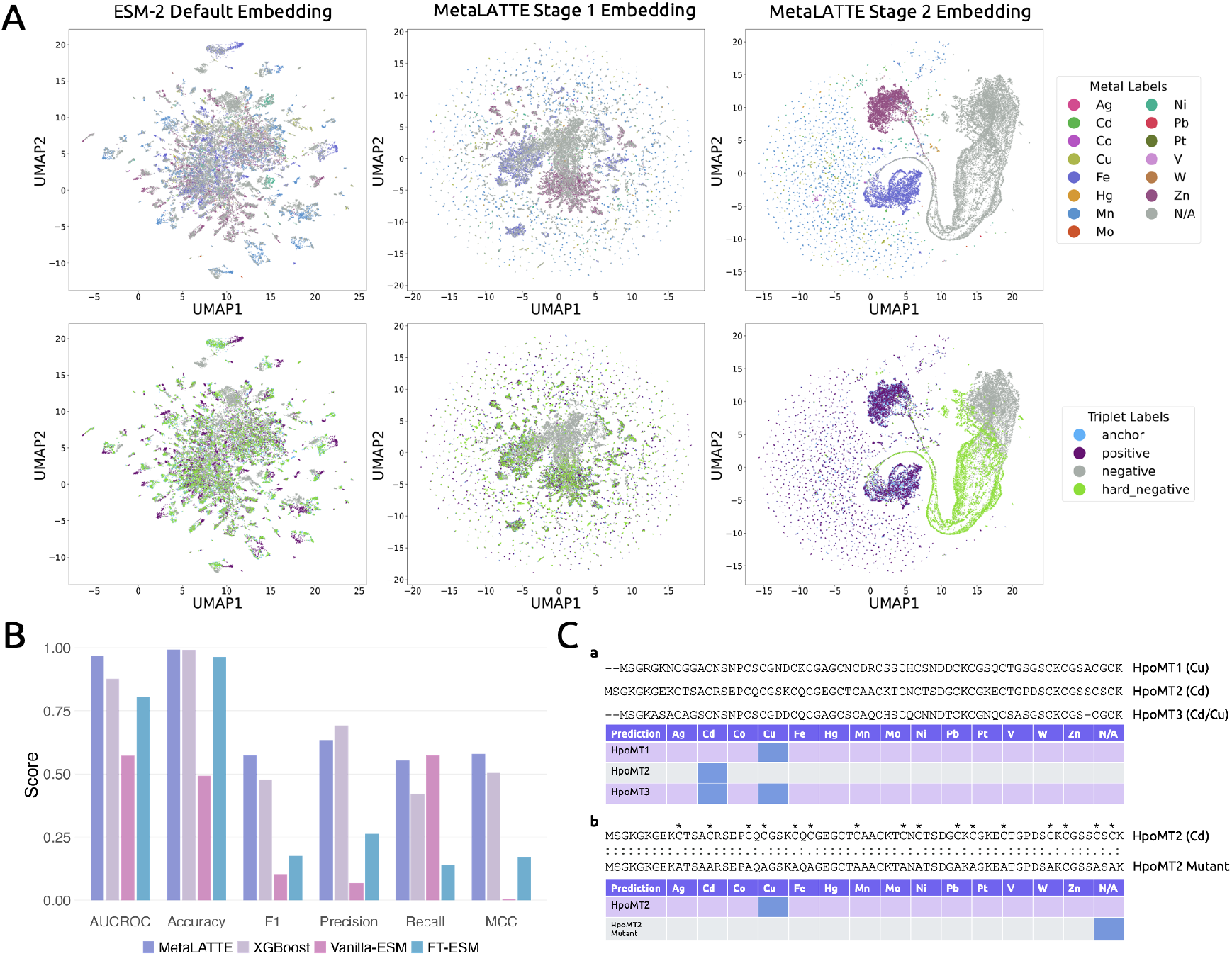
MetaLATTE captures label differences from the triplet loss. **A)** Protein embeddings from the validation set visualized in UMAP. Top row: colored by different metal classes; Bottom row: colored by different triplet categories. **B)** Classification metrics comparison between different models averaged over all the metal-binding protein classes. **C)** a: Apply MetaLATTE to isoforms sequenced from cDNAs; b: Apply MetaLATTE to manual alanine-swapped sequences. Alanine swaps are labeled with *.

To validate MetaLATTE’s real-world applicability, we tested on understudied mollusk proteins from *Helix pomatia* [Chabicovsky et al., 2003, Calatayud et al., 2021, Baroudi et al., 2020], which present a challenging case due to their high sequence similarity between isoforms yet distinct metal specificities. Remarkably, MetaLATTE successfully distinguished Cu-specific (HpoMT1), Cd-specific (HpoMT2), and dual-binding (HpoMT3) isoforms despite 75.4% sequence identity between some pairs, and correctly identified the loss-of-function when binding site residues were mutated to alanines (Figure 2C). These results demonstrate MetaLATTE’s superior ability to predict metal-binding properties of poorly-characterized proteins, highlighting its potential for discovering novel metal-binding proteins in understudied organisms and validating its readiness to guide Metalorian’s peptide generation process.

### Metalorian uses MetaLATTE-guided diffusion sampling to generate heavy metal binding peptides

While MetaLATTE enables accurate prediction of metal-binding potential from sequence alone, generating new peptides that bind specific metals, especially underrepresented or toxic ones, poses an even greater challenge [Luo et al., 2024]. Existing design approaches rely on fixed scaffolds, structure-based modeling, or directed evolution [Learte-Aymamí et al., 2024, Chabicovsky et al., 2003], all of which limit diversity, controllability, or feasibility for disordered motifs in natural metalloproteins. Moreover, structure-guided generation tools like AlphaFold3 often perform poorly on metal-binding peptides [Zhai et al., 2025], which are typically cysteine- or histidine-rich and disordered in the absence of a bound metal, resulting in low-confidence (low predicted Local Distance Difference Test (pLDDT)) predictions.

To address these limitations, we introduce Metalorian (Figure 3A), a co-evolving conditional diffusion model [Lee et al., 2023] that generates *de novo* metal-binding peptides directly in the latent space of protein embeddings. Metalorian couples two denoising processes: a continuous diffusion over ESM-2 embeddings from MetaLATTE, and a discrete diffusion over metal labels, with bidirectional conditioning between them at every timestep. The model is trained with contrastive triplet loss to ensure alignment between sequences and intended metal classes, while a length control mechanism enables generation of minimal, tunable scaffolds.

**Figure 3:**
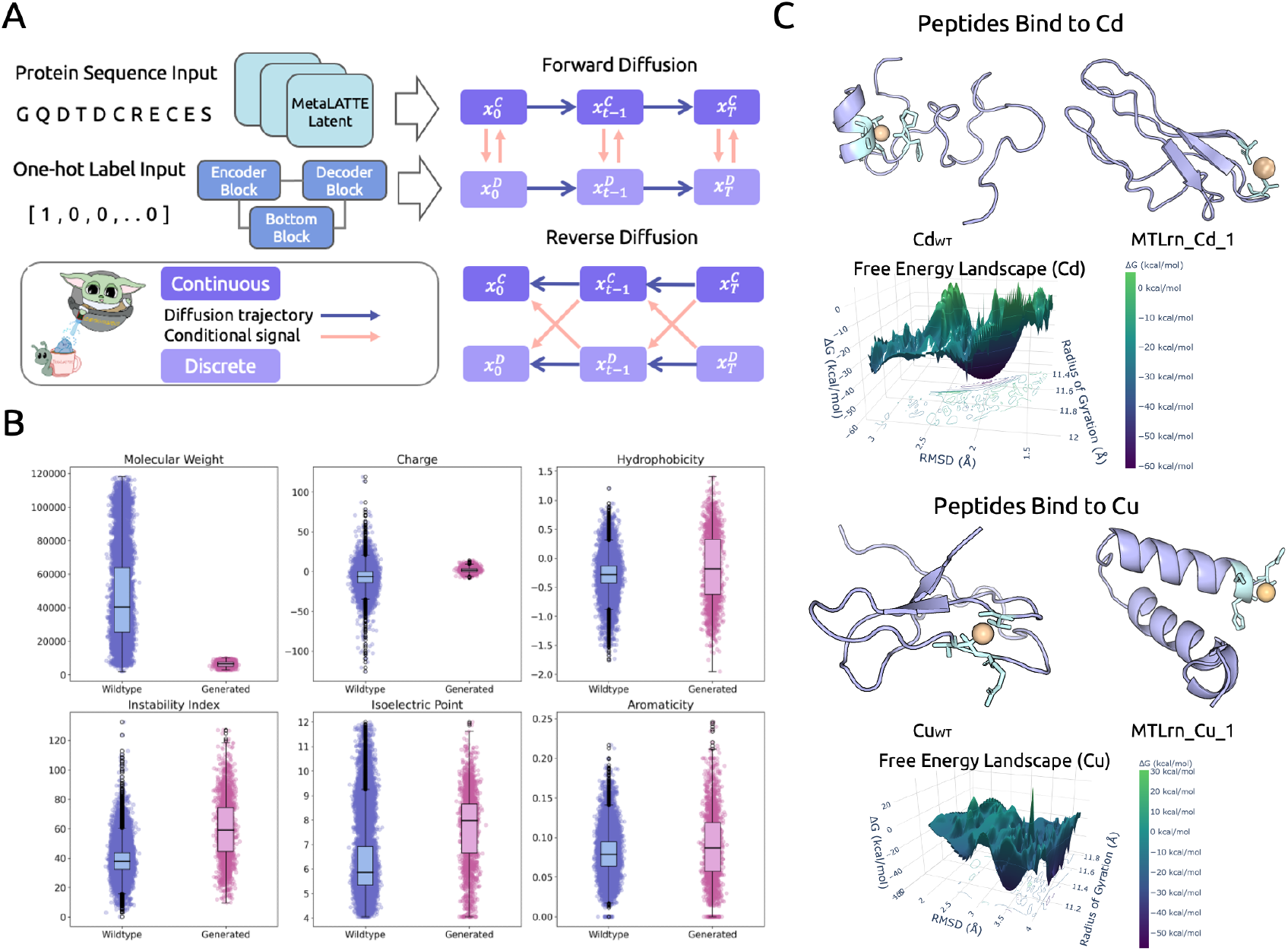
Metalorian schematic and analysis. **A)** Pipeline of the Metalorian architecture **B)** Biophysical property analysis between Metalorian-generated peptides vs. wild-type metal-binding proteins. **C)** Simulated selective Cd and Cu *de novo* peptides and free energy landscape analysis from corresponding MM/PBSA results.

Using our Metalorian diffusion model guided by the MetaLATTE classifier, we generated metal-binding peptide sequences with lengths between 30 and 80 residues for Cu, Zn, Cd, Co, and Ni. When compared to natural metal-binding proteins from the MbPA database [Li et al., 2023], our sequences demonstrated successful optimization with reduced molecular weight while preserving key features such as net charge distribution (Figure 3B). This controllable and reduced length and molecular weight (MW) is particularly significant, as lower-MW peptides have been associated with enhanced chelation activity [Xia et al., 2012, Luo et al., 2024], a key property needed for bioremediation.

Analysis of the generated peptides revealed several notable characteristics that align with known metalbinding principles. We observed an increase in the instability index in our designed peptides compared to their wild-type counterparts, which likely reflects a design bias toward short-lived, yet functionally competent, metal-chelating scaffolds. This structural behavior mimics documented metallothioneins, which are naturally occurring cysteine-rich peptides that lack defined structure in their apo state but adopt specific conformations only upon metal binding [Vašák and Hasler, 2000, Sutherland and Stillman, 2014, Ziller and Fraissinet-Tachet, 2018]. Additionally, the generated peptides exhibited increased hydrophobicity, isoelectric point, and aromaticity, driven by the enrichment of amino acids like cysteine, histidine, and phenylalanine (see Supplementary Data 4.1). Notably, cysteine and histidine residues contribute directly to metal binding through their electron-donating side chains. The thiol groups of cysteine and imidazole rings of histidine readily form coordinate bonds with transition metal ions such as Cu, Zn, and Cd, forming stable complexes [Permyakov, 2021, Mejáre and Bülow, 2001, Luo et al., 2024, Ziller and Fraissinet-Tachet, 2018].

Together, these features suggest that Metalorian captures key residue-level preferences of metal-binding proteins. The model optimizes toward minimal scaffolds analogous to metallothioneins, where cysteine and histidine residues dictate metal selectivity and affinity [Calatayud et al., 2021, Zhang et al., 2021].

### Metalorian-generated peptides exhibit potent binding capacities *in silico*

To validate the structural stability and metal-binding capability of our generated sequences, we performed detailed MD simulations comparing generated peptides with wild-type metal-binding proteins. Structural stability analysis revealed that across both Cu and Cd systems, generated peptides (MTLrn_Cu_1 and MTLrn_Cd_1) demonstrated comparable or improved structural stability relative to wild-types. Similar or lower backbone root-mean-square-deviation (RMSD) and radius of gyration values (Figure 3C; Figure S8-S9) indicated that the generated peptides maintain stable conformations during the simulation period. In addition, energetic decomposition from Molecular Mechanics Poisson–Boltzmann Surface Area (MM/PBSA) analysis revealed that the generated peptides achieve consistently exhibited stronger electrostatic interactions (ΔEEL) compared to wild-type peptides, which is particularly significant as electrostatic interactions are a key driver of metal coordination (Table S3) [Irving and Williams, 1953]. Despite the substantial solvation penalties that arise from desolvation effects in binding processes [Kolar et al., 2011], they maintained favorable net binding energies, indicating effective coordination *in silico*.

To further understand the conformational behavior of the generated peptides, we performed conformational landscape analysis via time-lagged independent component analysis (TICA). Both wild-type and generated peptides show similar patterns in their TICA projections, suggesting comparable conformational sampling during our simulation timescale. While longer simulations would be needed to definitively characterize the complete conformational landscapes [Schultze and Grubmuller, 2021], the similar TICA patterns indicate that generated and wild-type peptides exhibit comparable local conformational flexibility within the sampled timeframes (Figure S8C, Figure S9C).

### Metalorian-generated proteins bind Cu and Zn *in vitro*

To experimentally validate Metalorian-generated binders, we designed peptides against Zn and Cu, two metals that play vital roles as micronutrients but that can often exhibit cytotoxic effects at elevated levels [Van et al., 2024, Leal et al., 2018]. Candidate peptide binders were expressed as SUMO-tagged constructs in *E. coli*, purified via Ni-NTA affinity chromatography, biotinylated, and cleaved with TEV protease prior to analysis (Figure 4A). Using a novel sandwich ELISA based on a Cu- or Zn-coated microplate capture surface, we tested three *de novo* designed binders: MTLrn_Cu_1, MTLrn_Cu_2, and MTLrn_Zn_1 (MetaLATTE probabilities: 0.11, 0.82, 0.99, respectively), alongside a wild-type metalloprotein, Cu_WT_.

**Figure 4:**
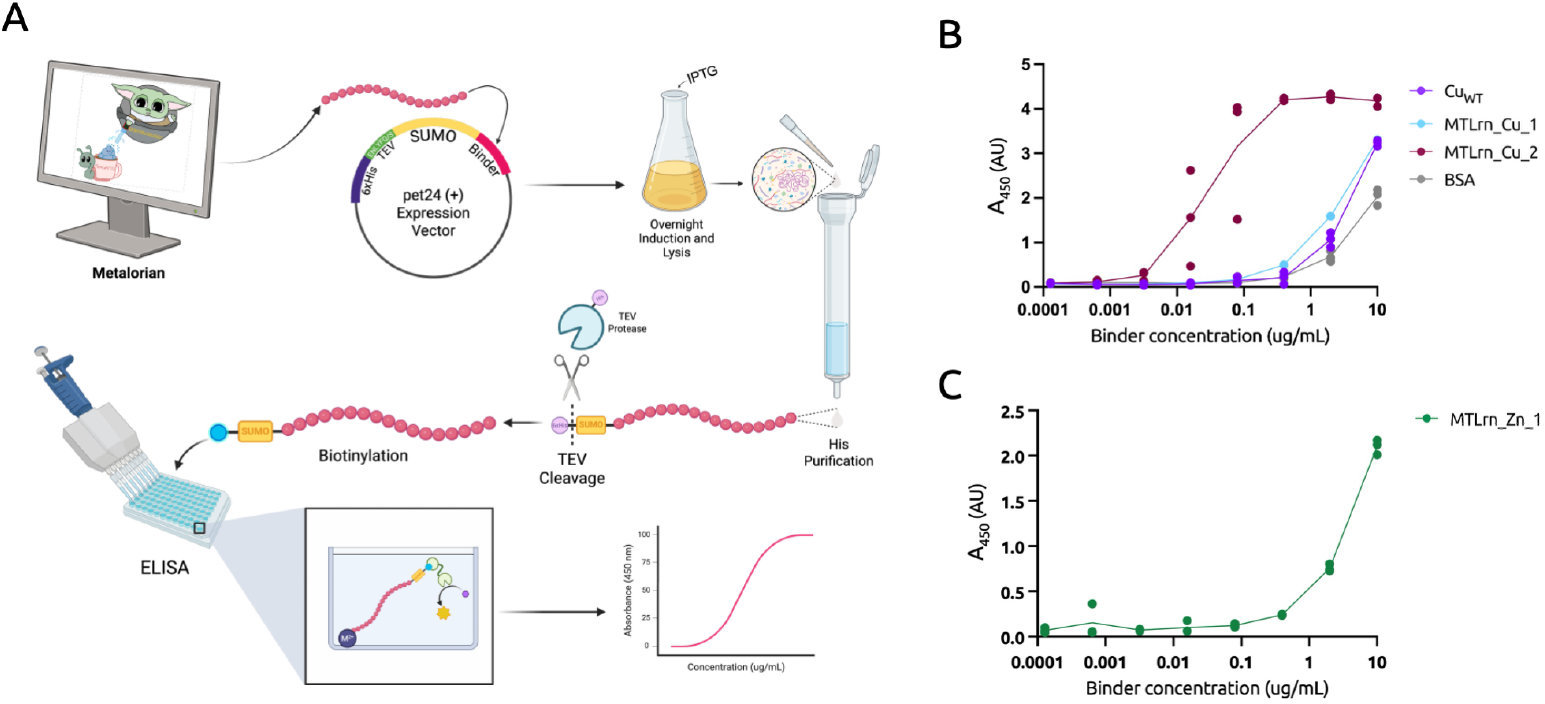
Experimental validation of Metalorian-generated binders *in vitro*. Two *de novo* Metaloriandesigned Cu binders, MTLrn_Cu_1 and MTLrn_Cu_2, were compared to a known Cu-binding protein, Cu_WT_, and BSA. (A) Experimental Screening pipeline, illustrating in-silico binder design, SUMO-fusion to increase solubility, and construct purification, cleavage, biotinylation, and ELISA-based analysis. (B) ELISA-based binding analysis of biotinylated SUMO-tagged constructs. Proteins were serially diluted in PBS and incubated with Cu-coated 96-well plates. Bound protein was detected via HRP-conjugated streptavidin and TMB substrate, with absorbance measured at 450 nm. (C) Binding of construct MTLrn_Zn_1 across a serial dilution series to Zn-coated plates. Technical triplicates are graphed at each dilution. Schematic made using BioRender.

Experimental results confirmed our computational predictions. MTLrn_Cu_1 exhibited a binding profile similar to Cu_WT_, with both showing higher absorbance relative to the BSA control (Figure 4B), consistent with their comparable behavior in MD simulations (Figure S8). MTLrn_Cu_2 demonstrated even stronger binding, with low-nanomolar affinity and a substantially improved Cu-binding profile compared to Cu_WT_, a well-characterized Cu metalloprotein (Figure 4B). Similarly, MTLrn_Zn_1 showed robust binding to Zn-coated microplates at mid-nanomolar concentrations (Figure 4C). Taken together, these results highlight the capability of Metalorian to generate novel metal-binding peptides that rival or exceed the affinity of known metalloproteins.

## Discussion

Heavy metal contamination represents a pressing global challenge, with regions characterized by intensive industrial operations, notably mining districts and densely populated urban areas, bearing the brunt of this environmental burden. This contamination affects nearly 40% of the Earth’s landmass [Tchounwou et al., 2012] and 14 billion people [Hou et al., 2025], creating an urgent need for effective remediation strategies. Heavy metal-binding peptides offer promising bioremediation tools for metal contamination cleanup [Luo et al., 2024], motivating our development of Metalorian, a novel peptide-design platform that addresses current limitations in the field. Leveraging CoDi, a co-evolutionary diffusion framework [Lee et al., 2023], in conjunction with MetaLATTE’s multi-label metal-binding guidance, the model systematically designs metal-binding peptides. By integrating continuous protein embeddings with discrete sequence properties, our approach systematically designs shorter and lighter metal-binding peptides enriched in the key residues for metal coordination.

Overall, the synergy between data-driven models and a robust experimental validation platform offers a scalable framework for designing and experimentally validating novel metal-binding peptides, with potential applications in environmental remediation. In the future, we aim to expand Metalorian’s scope to additional heavy metals and multiple binding scenarios while extending our experimental validation to additional candidates. The platform’s modular framework allows for straightforward adaptation to new target metals, off-target-avoiding binding contexts [Vincoff et al., 2025a, Vincoff et al., 2025b], and multi-objective property design [Tang et al., 2025, Chen et al., 2025], suggesting broad applicability across diverse remediation challenges. Finally, this study highlights a crucial connection between biodiversity conservation and technological innovation. Given the limited resources during training, we hope that this study also raises awareness about the need to protect biodiversity and invest in basic biology, as the solutions to current environmental problems may lie within understudied organisms yet to be discovered.

## Methods

### Dataset Preparation

#### Metal-Binding Protein Data Curation

A comprehensive dataset of metal-binding proteins was curated from multiple sources to support both our generation and classification objectives. The primary data source was the MbPA database [Li et al., 2023], from which proteins binding to transition and heavy metals are selected: silver (Ag), cadmium (Cd), cobalt (Co), copper (Cu), iron (Fe), mercury (Hg), manganese (Mn), molybdenum (Mo), nickel (Ni), lead (Pb), platinum (Pt), vanadium (V), tungsten (W), and zinc (Zn) (Figure 1A). To ensure sufficient representation, metal labels with fewer than 6 samples were excluded. Non-binding protein samples were obtained from the Mpbipred database [Aptekmann et al., 2022] to provide negative examples and create a balanced training set.

During benchmark evaluation, three different sources were used: (1) a stratified validation split from our stage 2 contaminated dataset, (2) a non-overlapping dataset from MetalPDB [Andreini et al., 2012] containing metal-binding proteins absent from our training data, and (3) newly sequenced mollusk genomic data [Calatayud et al., 2021] not documented in existing protein databases at the time of this study. It is important to note that certain transition metals (e.g., Pb, V) are underrepresented in the held-out test set due to limited study, with the test data predominantly featuring well-studied metal ions such as Zn and Mn.

#### MetaLATTE: Classification Model Data Processing

The classification model training employed a two-stage approach with progressively challenging negative examples:

##### Stage 1 Training

The model was initially trained on the basic distinction between different types of metalbinding proteins, using balanced stratification separation [Sechidis et al., 2011] for 4-fold cross-validation to assess model performance and generalization. Proteins from the Mpbipred database documented as non-metal binding and are labeled as N/A.

##### Stage 2 Training

Two tiers of negative examples were constructed to enhance site-specific learning S1:

- “Semi-hard negatives”: Decoy sequences created by modifying binding site amino acids in metalbinding proteins using BLOSUM62 matrix-guided substitutions [Henikoff and Henikoff, 1992], replacing residues within a ±3 window around binding sites (Figure 1B)
- “Hard negatives”: Sequences with alanine substitutions at the precise binding sites identified in the MbPA database

A triplet training approach [Schroff et al., 2015] was used to organize data into [anchor, positive, negative] triplets for triplet loss minimization. Each anchor sequence was paired with a positive sequence sharing the same metal-binding labels and a corresponding negative sequence. As training progressed, the batches gradually transitioned from easy to hard negatives, allowing the model to distinguish increasingly subtle functional differences.

To address the class imbalance, an oversampling strategy was applied for minority metal labels, generating multiple triplets for anchor sequences with rare binding profiles (e.g., Cd, V, W) (Supplementary Description 1).

#### Metalorian: Generation Model Data Processing

Single-property optimization was focused by selecting single-ion binding proteins from the curated dataset. A standard 80/20 train/validation stratification split was chosen to ensure robust model evaluation. Unlike the classification model training, the original, unaugmented dataset was used for the diffusion model to optimize computational efficiency while preserving the essential binding characteristics required for generation. This approach was necessary as the contaminated dataset from the classification pipeline would have tripled the data size, making diffusion model training computationally expensive. Importantly, Metalorian leverages the latent space representations learned by MetaLATTE, making the classification model an essential prerequisite for our diffusion-based generation approach.

#### Model Architecture

The base for the classifier was built on top of the pre-trained ESM-2-650M pLM [Lin et al., 2023], which has already acquired basic protein evolutionary information and is well-validated in the literature for both prediction and design tasks [Brixi et al., 2023, Bhat et al., 2025, Chen et al., 2023a, Su et al., 2023, Peng et al., 2025, Vincoff et al., 2025c].

#### MetaLATTE: Classification Model

MetaLATTE was designed to detect metal-binding properties within protein sequences, specifically focusing on the recognition of site-specific binding motifs that traditional protein language models might not sufficiently capture. ESM-2-650M was fine-tuned for our multi-label classification task, unfreezing the last two layers to adjust its pre-trained embeddings while preserving most of the foundational protein knowledge. To enhance site-specific sensitivity, an attention pooling layer and rotary position embeddings [Su et al., 2024] were applied before the classification head.

A two-stage training approach was applied. For stage one, a multi-label classification model was trained first across 14 heavy metal labels plus a non-binding label. The batch size was set to 6, with a peak learning rate of 1e-3 using a cosine learning rate scheduler with 200 warm-up steps [Touvron et al., 2023]. Classification thresholds were dynamically updated through exponential moving average (EMA) [Yan et al., 2022] with a historical memory factor (*α*) of 0.6 (Algorithm 1). The max epoch was set to 30 with early stopping based on validation macro-F1 scores. Training was completed within 32 GPU hours on a 7×NVIDIA A6000 DGX server with 350 GB of shared VRAM using PyTorch Lightning for distributed data-parallel training [Falcon and The PyTorch Lightning team, 2019]. For stage two, building on the Stage 1 model, triplet-based contrastive learning [Schroff et al., 2015, Balntas et al., 2016, Paszke et al., 2019, Liu et al., 2019] was applied to enhance discrimination between binding and non-binding sequences, particularly for challenging cases. A cosine-scheduled learning rate would peak at 2e-4 with 100 warm-up steps and EMA memory factor was increased to 0.9 for more stable threshold updates. The triplet margin parameters were carefully tuned to balance between hard and easy examples. Early stopping tracked validation macro-F1 scores, with optimal performance typically achieved around epoch 15. The same hardware setup completed this stage in 56 GPU hours, using dynamically pre-batched datasets to optimize training efficiency (Algorithm 3).

#### Loss Functions

MetaLATTE’s training incorporates four complementary loss functions: *Class-Balanced Focal Loss*: Addressed class imbalance [Cui et al., 2019] with hyperparameters *γ* = 1.2 and *β* = 0.9999:

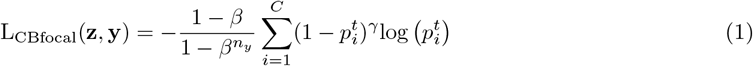

Where

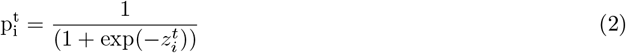

##### F1 Loss

Optimized the balance between precision and recall:

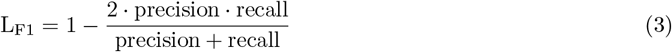

##### Reconstruction Loss

Preserved natural protein distribution patterns through cross-entropy. The reconstruction layers teach the model how to project embeddings back to logits with cross-entropy loss, enabling downstream sequence design:

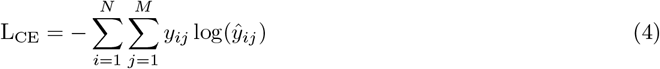

where *N* is the sequence length and *M* is the vocabulary size inherited from the ESM-2 model.

##### Triplet Loss (Stage 2 only)

Enhanced embedding space discrimination with adaptive margins [Liu et al., 2019] (Algorithm 2):

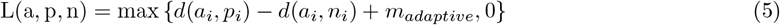

where *i* is the minibatch, and

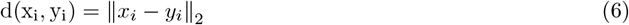

with adaptive margin:

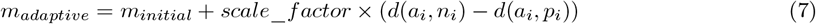

where *scale*_*factor* was set to 0.1. After epoch 10, semi-hard triplets were gradually replaced with hard triplets for refined embedding representation (Algorithm 4). The contribution of each loss function was tuned empirically. In Stage 1, focal, F1, and reconstruction losses were weighted equally. In Stage 2, triplet loss received higher weighting to prioritize embedding space refinement.

#### Metalorian: Generation Model

Metalorian adapts the co-evolving conditional diffusion framework [Lee et al., 2023] to protein sequence generation, leveraging the latent representations learned by MetaLATTE. The co-evolving training strategy integrates two coupled diffusion processes:

#### Continuous Protein Diffusion Model

Operates on MetaLATTE’s ESM-2 protein embeddings 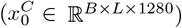 with the last 10 layers unfrozen. The forward process follows:

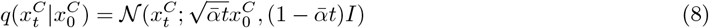

where 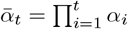 and *α*_*t*_ = 1 − *β*_*t*_.

The reverse process is parameterized by 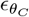 and conditioned on discrete variables:

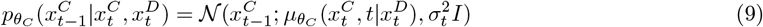

#### Discrete Label Diffusion Model

Operates on metal-binding labels 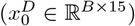 using a multinomial diffusion process with a TabularUnet backbone. The forward process uses categorical distributions:

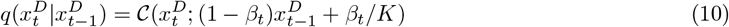

where *K* is the number of classes.

The reverse process is conditioned on continuous embeddings:

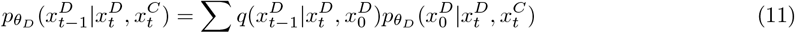

Metalorian was trained on 7×NVIDIA A100 GPUs with a batch size of 140 and a learning rate of 2 × 10^−4^ using the AdamW optimizer [Loshchilov and Hutter, 2019]. The implementation and parallelization utilized the PyTorch Lightning framework [Falcon and The PyTorch Lightning team, 2019]. During the generation phase, a sequence length control mechanism applied via masks allowed the generation of peptides within a controllable range, for example, 30 to 80 amino acids.

#### Loss Functions

The models were trained jointly with a combined loss function:

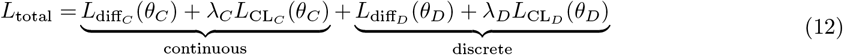

where 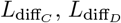 are diffusion losses for continuous and discrete components; 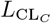, 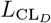 are contrastive learning losses with negative sampling; *λ*_*C*_, *λ*_*D*_ are weighting coefficients.

The contrastive learning loss uses a triplet formulation with positive and negative samples:

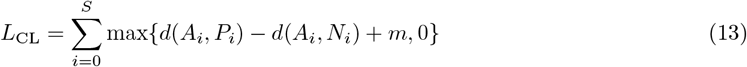

where *A* is the anchor, *P* is a positive sample, *N* is a negative sample, *d* is a distance metric, *m* is the margin, and *S* is the number of samples. This approach encourages the model to learn the true correlation between continuous embeddings and discrete labels while being robust to mismatched conditions.

For positive sample generation, 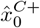 was generated conditioning on the matching discrete label 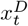 and 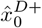 conditioning on the matching continuous embedding 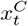. For negative sample generation, within a minibatch, 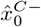 was generated using mismatched discrete label 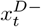 and 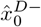 using mismatched continuous embedding 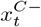. Additional implementation details, including visualizations of the co-evolving diffusion process and triplet sampling strategy, are provided in Figures S2–S3.

### Sampling

#### Progressive Verification Sampling

Two complementary sampling approaches for protein sequence generation would be used for metal-binding properties. The first approach, Progressive Verification Sampling (Algorithm 5), is designed for well-represented metal classes in our training data. During phase one (steps *T* to *T*_*c*_), it follows standard diffusion sampling:

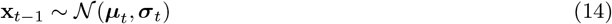

In phase two (steps *T*_*c*_ to 0), a verification mechanism was introduced that ensures label alignment through multiple sampling attempts. At each timestep, both predictor probability *P*(**x**_*t*_)[*y*_target_] from the MetaLATTE classification model and discrete label alignment 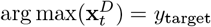 were evaluated. Sampling continues until success criteria are met: predictor confidence exceeds threshold *τ* and labels align. At each timestep, the continuous and discrete models inform each other’s sampling process:

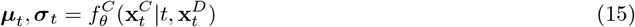

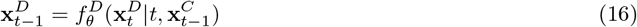

where 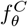 conditions the continuous sampling on the current discrete state, and 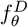 updates the discrete state based on both previous discrete state and new continuous sample.

#### Gradient-Guided Sampling

For metal classes with limited training examples or complex binding patterns, the Gradient-Guided Label Sampling (Algorithm 6) was applied. This approach extends classifier guidance [Dhariwal and Nichol, 2021] with dynamic scaling [Dinh et al., 2024] and label alignment. At each timestep *t*, we compute:

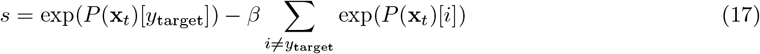

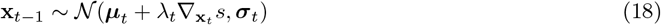

where *λ*_*t*_ doubles the base guidance scale *λ* when the predictor’s confidence for the target class falls below threshold *τ*. This co-evolution of continuous and discrete states, combined with gradient guidance, ensures that both sequence generation and label prediction remain consistent throughout the sampling process. The key difference here is that in gradient-guided sampling, this cross-model interaction works alongside the gradient guidance, while in progressive verification sampling it works with the verification mechanism.

#### Benchmarks Performance Metrics

For multi-label classification results, `scikit-learn` (v1.3.2) [Pedregosa et al., 2011] was used to compute sensitivity (recall), precision, macro F1-score, Matthews Correlation Coefficient (MCC), Area Under the Receiver Operating Characteristic Curve (AUROC), and proportional accuracy score. For the mollusk sequence analysis, sequence alignment was performed with LALIGN from EMBL-EBI [Pearson, 1991] and Geneious alignment, both utilizing the BLOSUM62 matrix [Henikoff and Henikoff, 1992], to establish sequence similarity relationships. For the sequences generated from Metalorian, biophysical properties were computed with the `Biopython` (v1.78) `ProteinAnalysis` [Cock et al., 2009], focusing on key attributes including: overall charge, hydrophobicity (Kyte-Doolittle Hydrophobicity scale [Kyte and Doolittle, 1982]), molecular weight, and instability index. The metal-binding propensity of generated sequences was evaluated using our pre-trained MetaLATTE classifier.

#### Benchmark Models

XGBoost [Chen and Guestrin, 2016], a state-of-the-art gradient boosting algorithm known for its strong performance on high-dimensional data, was chosen as the benchmark model. The XGBoost model was rigorously optimized using `optuna` [Akiba et al., 2019] for hyperparameter tuning, with the best model obtained after 200 trials. To ensure fair comparison, this optimized model was trained on ESM-2 default embeddings and incorporated the same EMA threshold update mechanism used in MetaLATTE. Two additional ESM baseline models were also trained. A vanilla ESM model (zero-shot) baseline used the Hugging Face [Wolf et al., 2019] `EsmForSequenceClassification` model without any fine-tuning, relying solely on pretrained knowledge. A head-finetuned ESM baseline used the same ESM-2 backbone, but only the classification head was trained while keeping the transformer layers frozen. All models were trained on identical training datasets and evaluated using the same validation splits to ensure consistent and fair comparison.

#### Embedding Visualization

UMAP with Louvain clustering from `scanpy` (v1.9.8) [Wolf et al., 2018] was used to visualize the shifting of learned embeddings space for the validation dataset after MetaLATTE training. The number of neighbors parameter in UMAP was set to 50.

#### Classical Molecular Dynamics

Structural models of peptide-metal complexes were first generated with AlphaFold3 (2024.08.19 update) [Abramson et al., 2024]. For consistent metal ion placement, docking was performed using Autodock VINA [Trott and Olson, 2010]. System preparation followed the AmberTools24 [Case et al., 2023] pipeline, using MCPB.py [Li and Merz Jr, 2016] to define bonded and nonbonded parameters, and GAMESS-US [Barca et al., 2020] for quantum mechanical optimization and RESP charge derivation of the metal binding site. Detailed rationale for this hybrid modeling approach is discussed in Supplementary Section 3.2.

For most transition metals such as Cu, the 6-31G(d, p)/LANL2TZ basis set [Roy et al., 2008] was used. For Cd, however, as the LANL2TZ basis set was insufficient due to Cd’s larger atomic radius, SBKJC-ECP basis set [Stevens et al., 1992] was chosen with its tighter convergence criteria.

The peptides were solvated with TIP3P water in a periodic box with a minimum distance of 10Å between the peptide and box edge. The system was neutralized with Na+ and Clions at a concentration of 0.155 mM to mimic the experimental condition of physiological conditions (PBS). The AMBER19SB force field [Tian et al., 2019] was used to represent amino acids, while the GAFF force field [Wang et al., 2004] was employed for the remaining atoms. Amber preparation to GROMACS transition was performed using `acpype` (v2023.10.27) [Sousa da Silva and Vranken, 2012].

Following the standard GROMACS 5.0 [Abraham et al., 2015] workflow, energy minimization (EM) was first performed using the steepest descent algorithm for 50,000 steps with an energy step size of 0.01 and a convergence criterion of 1000.0 kJ/mol/nm. Position restraints were applied to maintain structural integrity during minimization. After minimization, the NVT ensemble was performed for temperature equilibration with no pressure coupling (pcoupl = no). The system was directly heated to 300K and equilibrated for 100 ps (50,000 steps with a 0.002 ps time step) under position restraints. Temperature coupling was handled using the V-rescale thermostat [Bussi et al., 2007] with protein and non-protein atoms in separate coupling groups (*τ* = 0.1 ps). Velocity generation was enabled with a target temperature of 300K. Pressure equilibration in the NPT ensemble was conducted using the c-rescale barostat [Bernetti and Bussi, 2020] at 1 bar with *τ* = 2.0 ps and a compressibility of 4.5 × 10^−5^bar^−1^. The NPT equilibration ran for 100 ps (50,000 steps with a 0.001 ps time step) under continued position restraints. Temperature was maintained at 300K using the V-rescale thermostat with the same coupling parameters as the NVT phase.

Production MD simulations were performed using stochastic dynamics integrator for 2.5 ns (5,000,000 steps with a 0.0005 ps time step), with position restraints being removed. Temperature was maintained at 300K using V-rescale thermostat, and pressure was controlled at 1 bar using c-rescale barostat with *τ* = 2.0 ps. Long-range electrostatic interactions were handled using the Particle Mesh Ewald (PME) method [Petersen, 1995] with a cutoff of 1.2 nm. All bonds were constrained using the LINCS algorithm [Hess et al., 1997]. Coordinates, velocities, and energies were saved every 6.25 ps for subsequent analysis.

Trajectory analysis was conducted using `CPPTRAJ` [Roe and Cheatham III, 2013], and binding stability was assessed through free energy calculations using `gmx_MMPBSA` [Kollman et al., 2000]. Root-mean-square deviation (RMSD) was calculated for peptide backbone and binding site residues to assess structural stability. Root-mean-square fluctuation (RMSF) was calculated to identify flexible regions. Radius of gyration (Rg) was computed to monitor the compactness. Time-lagged independent component analysis (TICA) was performed using `PyEMMA` [Scherer et al., 2015] with a lag time of 10 frames. Final structural visualization was performed using `PyMOL` [Schrödinger, LLC, 2015] (v 3.1), where residues with polar contacts closer than 3.0Å to the metal ion or coordinating residues were identified and annotated.

#### Generation and Expression of SUMO Fusion Constructs with Modified TEV Sites

The expression vector containing an N-terminal 6xHis-SUMO tag was derived from the construct described in [Chen et al., 2023a] and modified to reposition the TEV protease cleavage site upstream of the SUMO tag. To do so, the vector was linearized via restriction digestion with NdeI (New England Biolabs (NEB), R0111S) and BamHI-HF (NEB, R3136S), and assembled using Gibson Assembly (NEB, E2621L) with double-stranded DNA gBlocks (Integrated DNA Technologies) encoding the modified sequences. Sequences of assembled constructs were verified via Sanger sequencing (Genewiz) and subsequently transformed into chemically competent *Escheria coli* BL21 (DE3) cells (NEB, C2527H). Two-milliliter starter cultures were picked from a glycerol stock and grown overnight at 37°C with shaking in 2xYT (Sigma Aldrich, Y2377) supplemented with 100 µg mL^−1^ kanamycin. Starter cultures were then diluted 1:500 into bulk cultures and grown to an optical density (OD_600_) of 0.6–0.8 before induction with 1 mM isopropyl ***β***-D-thiogalactopyranoside (IPTG) and subsequent overnight incubation at 37°C with shaking (225 RPM). Cells were collected by centrifugation (4,500 x g) at 4°C and washed twice with ice-cold 1x PBS. The resulting cell pellets were frozen at −20°C overnight, thawed to room temperature, then lysed using BugBuster protein extraction reagent (Millipore Sigma, 70584-3) supplemented with recombinant lysozyme (Millipore Sigma, 71110-3) and benzonase endonuclease (Millipore Sigma, E1014-25KU) for 20 minutes at room temperature with gentle rocking. The lysate was diluted with lysis buffer (1x PBS, 20 mM imidazole, 300 mM NaCl, 1x Halt protease inhibitor cocktail (ThermoFisher, 78430), pH 7.4), then centrifuged at 14,000 x g for 30 minutes. The cleared supernatant was mixed end-over-end at 4°C for 1 hour with HisPur Ni-NTA resin (ThermoFisher, 88221) equilibrated with 20 mM imidazole and 300 mM NaCl in 1x PBS. The resin was collected by centrifugation at 700 x g for 2 minutes, washed twice with 25 mM imidazole in 1x PBS, and once with 50 mM imidazole. Bound proteins were eluted with three sequential washes using 500 mM imidazole. Eluates were concentrated using Amicon Ultra centrifugal filters (3K MWCO; Millipore Sigma, UFC900308) and desalted with Zeba Spin Desalting Columns (7K MWCO; ThermoFisher Scientific, 89892). Expression and purity of proteins in soluble, insoluble, and purified fractions were assessed by SDS–PAGE. Protein concentrations were determined using the Qubit Protein Assay Kit (ThermoFisher, Q33211).

#### Enzymatic Cleavage and Biotinylation of SUMO-Tagged Peptides

To remove the N-terminal 6xHis tag, constructs were cleaved using TEV protease (NEB, P8112S) following the manufacturer’s protocol, scaled for 100 µg of protein and incubated at 30°C for 2 hours, then subjected to a reverse His-purification. The entire cleavage solution was mixed end-over-end at 4°C for 1 hour with HisPur Ni-NTA resin equilibrated with 20 mM imidazole and 300 mM NaCl in 1x PBS. Resin was collected by centrifugation at 700 x g for 2 minutes, washed once with 50 mM imidazole in 1x PBS, and eluted with three consecutive washes using 500 mM imidazole. Cleaved and uncleaved fractions were analyzed by SDS–PAGE (Figure S7A), and cleavage was confirmed by anti-His tag Western blotting (Figures S6 and S7B). Flow-through fractions containing cleaved, His-tag-free protein were concentrated and buffer exchanged using 2 mL Zeba™ Spin Desalting Columns with a 7 kDa molecular weight cutoff (Thermo Fisher Scientific, 89877). Proteins and BSA were then biotinylated using the EZ-Link™ Micro Sulfo-NHS-Biotinylation Kit (Thermo Fisher Scientific, 21925) according to the manufacturer’s instructions. Excess unreacted biotin was removed via a second round of desalting.

#### Anti-His Western Blotting

Cleaved and uncleaved purified constructs, as well as eluted fractions from reverse His-purification, were quantified, and 2 µg of each protein was mixed with 4X Bolt™ LDS Sample Buffer (Thermo Fisher Scientific) containing 5% *β*-mercaptoethanol. Samples were denatured by incubation at 95°C for 10 minutes, and 20 µL of each was loaded onto a Novex™ 4–20% Tris-Glycine Mini Protein Gel (Thermo Fisher Scientific, XP04205BOX) and separated by electrophoresis. iBlot™ 2 Transfer Stacks (ThermoFisher Scientific, IB24002) were used for transfer onto Polyvinylidene fluoride (PVDF) membrane. Following a 1-hour room-temperature incubation in 5% nonfat dry milk in 1x Tris-buffered saline (50 mM Tris-HCl, 150 mM NaCl) supplemented with 0.05% Tween-20 (v/v) (TBS-T), proteins were probed with an HRP-conjugated 6xHis Tag Monoclonal Antibody (Thermo Fisher Scientific, MA1-21315-HRP) diluted 1:5000 in 5% nonfat dry milk in 1x TBS-T for 1 hour at room temperature. Following three washes with 1x TBS-T for 5 minutes each, blots were detected by chemiluminescence using an Invitrogen iBright™ CL1500 Imaging System (Figures S6 and S7B).

#### Sandwich ELISA

Cu- and Zn-coated microplates (bioWORLD, 20140021) were blocked with 200 µL per well of SuperBlock Blocking Buffer in PBS (Thermo Fisher Scientific, 37515) according to the manufacturer’s instructions. Biotinylated SUMO-tagged constructs or biotinylated BSA control were prepared at 10 µg mL^−1^ in PBS and serially diluted in triplicate. A total of 200 µL of each dilution was added to the plates and incubated for 1.5 hours at room temperature with gentle shaking. Following incubation, plates were washed three times with PBS supplemented with 0.05% Tween-20 (PBS-T). HRP-conjugated streptavidin (Thermo Fisher Scientific, N100) was diluted 1:20,000 in PBS-T containing 1% bovine serum albumin (BSA, v/v) and added at 100 µL per well. Plates were then incubated for 1 hour at room temperature with gentle agitation, followed by three additional washes with PBS-T. Detection was carried out using 100 µL per well of 3,3’,5,5’tetramethylbenzidine substrate(1-Step Ultra TMB-ELISA; Thermo Fisher Scientific, 34029) for 15 minutes at room temperature, protected from light. The reaction was then quenched with 100 µL of 1 N HCl, and absorbance was immediately measured at 450 nm using a GloMax^®^ Discover plate reader (Promega). IC_50_ values were calculated using a sigmoidal four-parameter least-squares fit in GraphPad Prism version 10.

## Declarations

## Acknowledgements

We thank Mark III Systems and the Duke Computing Cluster for computing support. We further thank Willi Yu, Jing Guo, Kunal Mishra, and Tianlai Chen for their insights related to the manuscript. The work was funded by a Garden Grant from the Homeworld Collective to the lab of P.C.

## Author Contributions

Y.Z. designed, implemented, and validated MetaLATTE and Metalorian architectures *in silico*. D.S. designed and performed all experimental assays, with assistance from Z.Q. All authors wrote and reviewed the paper. P.C. conceived, designed, directed, and supervised the study.

## Data and Materials Availability

All data needed to evaluate the conclusions are presented in the paper and tables. MetaLATTE model weights and code are located at https://huggingface.co/ChatterjeeLab/MetaLATTE. An easy-to-use inference pipeline for MetaLATTE can be found at: https://huggingface.co/spaces/ChatterjeeLab/MetaLATTE-demo. Metalorian model weights and code are located at https://huggingface.co/ChatterjeeLab/Metalorian.

## Competing Interests

P.C. is a co-founder of Gameto, Inc., UbiquiTx, Inc., and AtomBioworks, Inc. and advises companies involved in peptide development. P.C.’s interests are reviewed and managed by the University of Pennsylvania in accordance with their conflict-of-interest policies. Y.Z., D.S., and Z.Q. have no conflicts of interest to declare.

## Supplementary Information

### 1 Data processing

#### The Metal-binding Protein Atlas (MbPA)

The original MbPA database contains 57,774 entries covering a wide range of species, including animals, plants, fungi, and microbes. It draws on protein sequences from the Protein Data Bank, UniProt, and AlphaFold DB. Residues within 3 Å of a metal ion in PDB structures are labeled as metal-binding sites [Li et al., 2023]. Since transition metals often display strong biases toward certain amino acids (compared to alkali or alkaline earth metals), we excluded entries that bind to non-transition metals [Li et al., 2023]. We excluded entire sequences rather than selectively removing non-transition metal annotations, as metal coordination, especially for transition metals, is known to induce conformational changes that may affect the overall structure [Vašák and Hasler, 2000, Simler et al., 2004]. Retaining only partial binding information may lead to inconsistencies in the residue-level context of coordination. We further removed sequences associated with cofactors such as heme, aiming to focus on proteins with intrinsic chelation ability rather than enzymatic activity driven by cofactors. Additionally, to mitigate potential crystallization artifacts, we excluded entries that reported only a single metal-binding site, which might not capture complete or biologically relevant binding environments.

After these filters, 46,765 entries remained. We then removed duplicate or single-binder sequences, arriving at 42,631 unique sequences with 50,374 metal labels in total. To increase the representation of less frequent metal labels (excluding zinc (Zn) and iron (Fe), negatives, and multi-label entries), we applied an oversampling factor of 5. Oversampling was performed by selecting the anchor and drawing 5 random positive sequences with the same label. Multi-labeled entries were excluded from oversampling due to their natural abundance and promiscuity.

We also collected 5,230 “easy negative” sequences (i.e., proteins without any metal-binding annotations and composed of only standard amino acids) [Aptekmann et al., 2022]. These easy negatives were used solely in the first-stage classification model for predicting “non-binding” (N/A) labels. They were not included during triplet-based training, which constitutes the second stage of the framework. After applying BLOSUM swaps, alanine scanning, and oversampling across batches, we finalized a dataset of 50,694 training and 12,729 validation pairs. All sequences were pre-batched with a maximum of 45,000 tokens per GPU for training (Algorithm 3).

#### Data Augmentation and Triplet Construction

As shown in Figure S1A, for single-labeled anchors, a positive was randomly drawn from sequences with the same metal label. For multi-labeled anchors, the positive was required to share at least two metal labels. Negative sequences were generated through residue corruption: BLOSUM-based substitutions produced semi-hard negatives, while alanine swaps yielded hard negatives by targeting residues at or near the binding site (Figure S1C). The train/validation split was performed using `MultilabelStratifiedShuffleSplit` [Sechidis et al., 2011] to preserve label distributions across splits.

To validate that the corrupted sequences remained biophysically realistic, we compared the charge and hydrophobicity distributions of corrupted versus natural sequences in the training set (Figure S1B). Given the large sample size, we performed effect size analysis between these two groups. Cohen’s *d* values were 0.094 for charge and -0.119 for hydrophobicity, both well below the conventional 0.2 threshold for small effects. Additionally, the high overlap coefficients (96.3% for charge and 95.3% for hydrophobicity) indicate that the distributions are highly similar. These findings suggest that while the corrupted sequences are biophysically realistic, metal-binding properties are subtle and difficult to distinguish based on global physicochemical features alone.

#### Manual alanine swap for benchmarking

For the isoform comparison, we applied alanine scanning mutagenesis to cysteine residues in HPOMT2, a Cd-binding protein isoform [Calatayud et al., 2021]. Cd coordination is strongly associated with thiol-rich environments, and cysteine residues are frequently the sole ligands in metallothionein-like Cd-binding motifs [Permyakov, 2021, Calatayud et al., 2021, Ziller and Fraissinet-Tachet, 2018]. Since direct annotation of coordinating residues was unavailable, we conservatively mutated multiple cysteines to alanines as a loss-of-function test to evaluate if our model had captured the residue-level metal-binding features.

**Figure S1:**
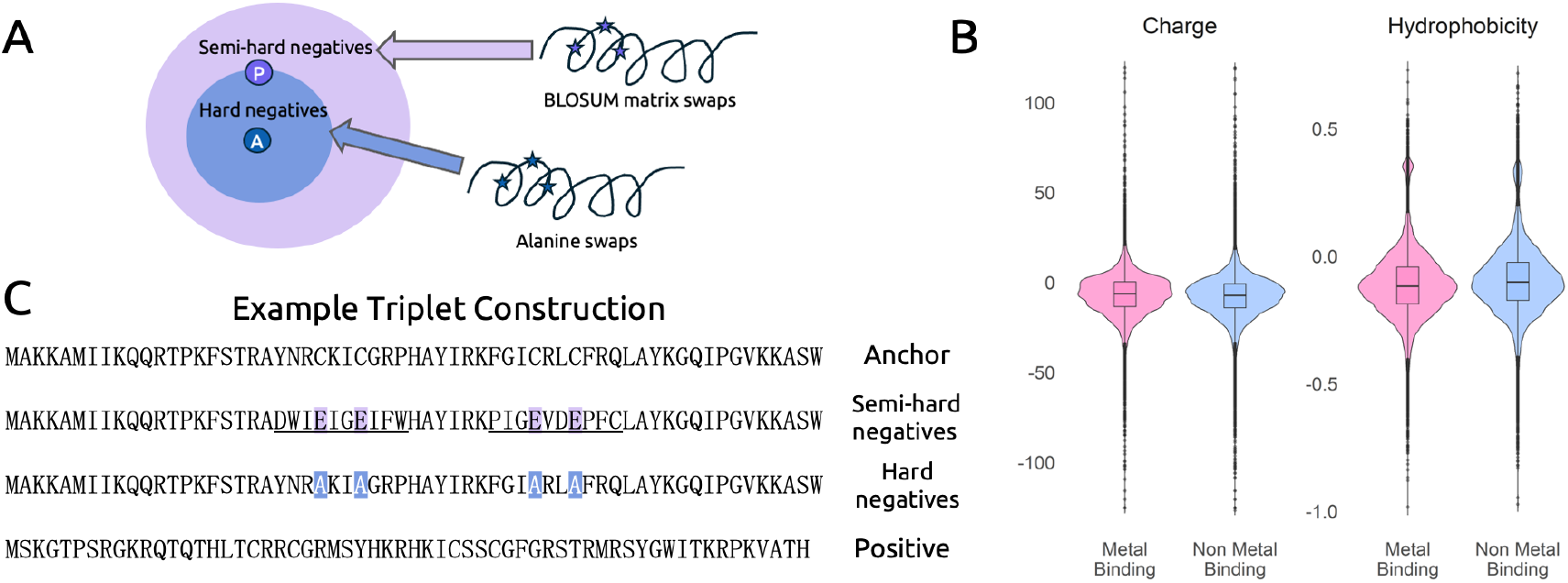
Illustration of data augmentation. A) Schematic triplet margins according to different labeling. Distances are shown arbitrarily here. B) Charge and Hydrophobicity distribution of the corrupted sequences compared to natural sequences. C) An example triplet pair of a Zn-binding protein. Anchor and Positive sequences are from the MbPA databases that share the Zn-binding label. Colored letters indicate binding sites got corrupted. Underlined residues in the semi-hard negative entries are other corrupted residues spanning the binding site in a window ±3 residues.

## 2 Algorithm explanation

**Figure S2:**
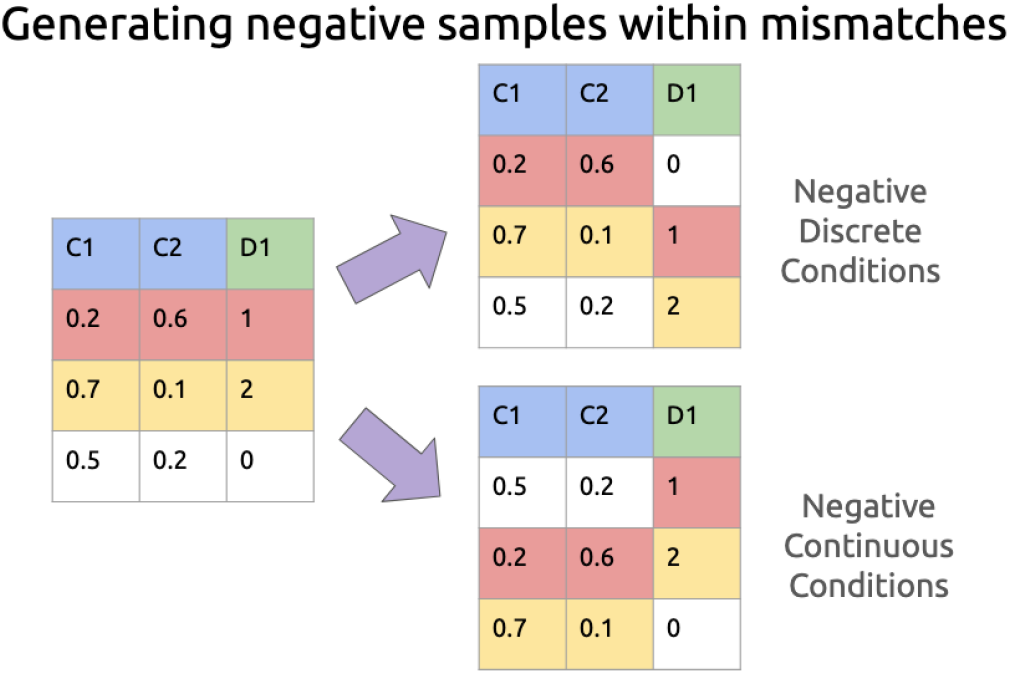
Triplet sampling for contrastive learning in Metalorian. Triplets are constructed using MetaLATTE latent embeddings 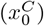 and metal-binding labels 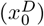. For continuous contrastive loss, a positive 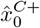 is generated from the matching 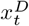, while a negative 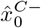 is generated from a mismatched label 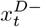 within the minibatch. An analogous process applies to discrete triplets 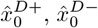, conditioned on continuous embeddings. This formulation encourages the model to bind continuous and discrete representations meaningfully across timesteps.

**Figure S3:**
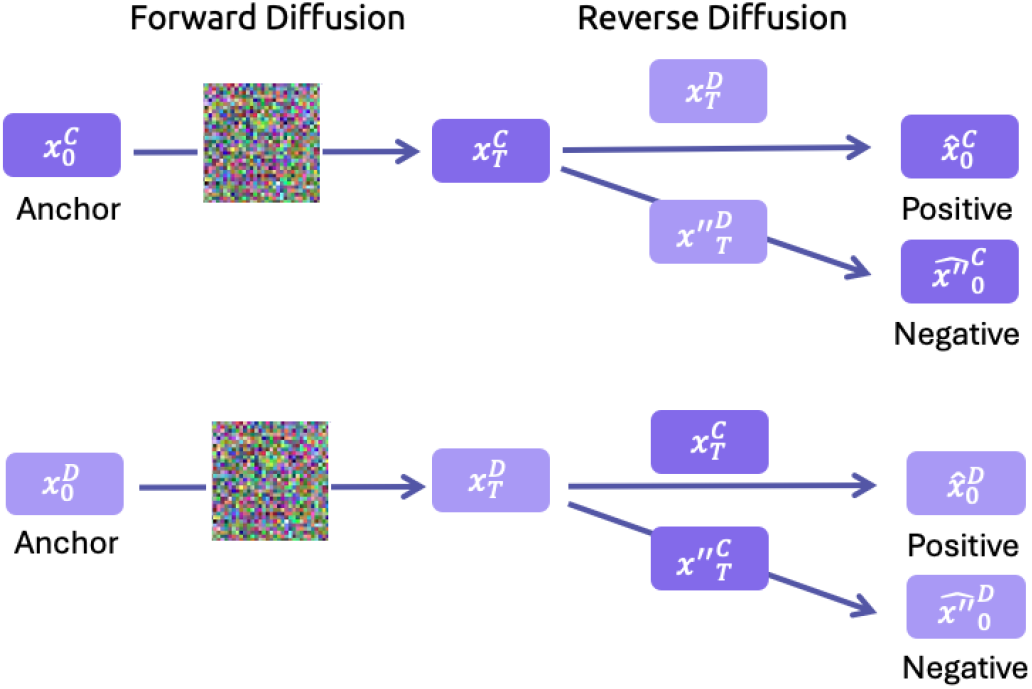
Reverse diffusion conditioning mechanism in Metalorian. At each denoising step *t* → *t* − 1, the continuous model predicts 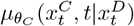 to obtain 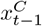, while the discrete model predicts 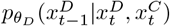. These coupled reverse processes enable co-evolving sampling, where each modality dynamically incorporates the updated state of the other.

## 3 Implementation Details

### 3.1 Model Architectures and Training Details

**Table S1:** Co-evolving Conditional Diffusion Model Architecture.

| Hyperparameter | Value |
| --- | --- |
| ESM Model Base | ESM2_t33_650M_UR50D |
| Max Sequence Length | 1280 |
| Training Batch Size | 140 |
| Diffusion Steps (T) | 50 |
| **Continuous Model (DiffusionProteinModel)** |  |
| Input Dimension | 1280 |
| Output Dimension | 1280 |
| Time Embedding Dimension | 1280 |
| Condition Projection | 15 → 1280 |
| Output Projection | 1280 |
| **Discrete Model (TabularUnet)** |  |
| Input Dimension | 15 |
| Condition Projection | 1280 → 15 |
| Output Dimension | 15 |
| Encoder Dimensions | [64, 128, 256] |
| Decoder Dimensions | [256, 128, 64] |
| Output Layer | 64 → 15 |

### 3.2 Structural Modeling Strategy and Rationale

Very limited options exist for *de novo* protein folding and binding prediction, especially across the diverse range of transition metals targeted in this study. To address this, we implemented a hybrid pipeline that combines AlphaFold3 (AF3) [Abramson et al., 2024] for initial structure prediction, AutoDock Vina [Trott and Olson, 2010] for binding pose sampling, and MCPB.py with GAMESS [Li and Merz Jr, 2016, Barca et al., 2020] for downstream quantum-classical geometry refinement and force field parameterization.

Although AF3 has demonstrated exceptional accuracy in modeling static protein-ligand complexes, its architecture does not explicitly consider electrostatics or coordination chemistry—both critical for transition metal binding [Zheng and Wang, 2025]. Similarly, AutoDock Vina lacks electrostatic and directional metal–ligand potentials [Sciortino et al., 2019], but employs Gaussian-shaped steric and hydrophobic terms alongside Monte Carlo optimization [Trott and Olson, 2010], enabling efficient sampling of initial poses in the absence of explicit coordination constraints.

Rather than using predefined grid potentials tailored to specific metal ions, which were only available for a limited subset such as Zn [Santos-Martins et al., 2014], we generated poses near the intended metal site and relied on downstream quantum mechanical optimization of GAMESS [Barca et al., 2020] for the corrections of coordination geometries and electronic properties.

The convergence of geometry optimization, MD trajectories, and MM/PBSA free energy calculations further supports the viability of our designed peptides. Overall, this pipeline bridges the limitations of current AI-driven folding and heuristic docking tools with rigorous quantum-classical refinement, enabling robust modeling of transition metal-binding peptides.

## 4 Extended Results

### 4.1 Amino Acid Composition Analysis

We observed a notable enrichment of histidine (H) and cysteine (C) in our generated metalloproteins, which aligns with their well-known roles in metal coordination [Lukács et al., 2021, Li et al., 2023] (Figure S4). Other enriched amino acids, such as proline (P) and glutamine (Q), may influence local conformation or solubility, potentially contributing to the overall structural preferences of the generated peptides [MacArthur and Thornton, 1991]. Conversely, residues like alanine (A), leucine (L), glycine (G), and valine (V), which are highly abundant across natural metalloproteins or proteins in general [uni, 2025] yet relatively less common at actual metal-binding sites, were depleted in our generated sequences. Overall the pattern demonstrates that our design captures the chemical preferences of the metal-binding environment.

A per-metal breakdown shows that Zn-binding designs strongly favor both cysteine and histidine, while Cd designs rely heavily on cysteine (Figure S5) [Ireland and Martin, 2019, Permyakov, 2021, Calatayud et al., 2021]. Meanwhile, nickel (Ni) and copper (Cu) designs are enriched in histidine, a residue frequently observed in native Ni/Cu binding motifs. Enrichment of phenylalanine (F), tyrosine (Y), and proline (P) may reflect their role in local conformational structuring, such as *π*–*π* stacking [Carter-Fenk et al., 2023]. Indeed, plant metallothioneins are known to feature aromatic-rich linker regions that may contribute to folding or domain organization [Ziller and Fraissinet-Tachet, 2018]. We also observed a preferential selection of histidine and cysteine (H/C) over acidic residues (D/E), consistent with the higher metal-binding affinity of imidazole and thiolate groups compared to carboxylates [Permyakov, 2021, Ziller and Fraissinet-Tachet, 2018].

**Table S2:**
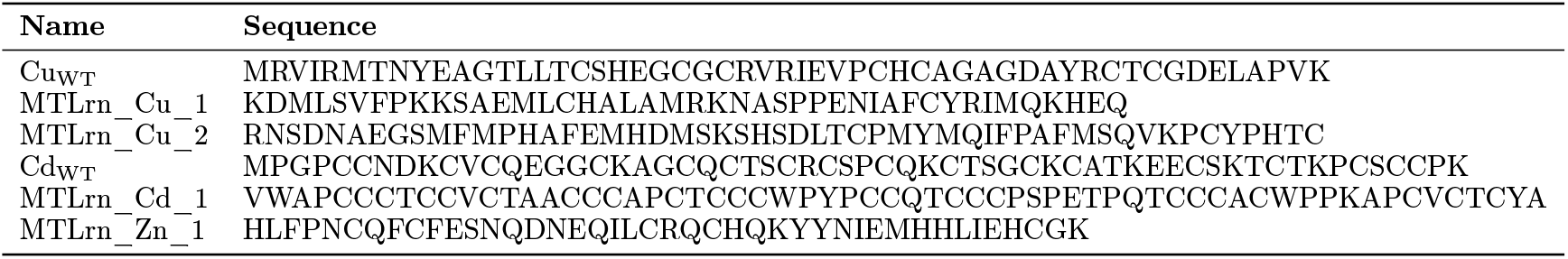
Sequences used for *in silico* modeling and *in vitro* validation.

**Figure S4:**
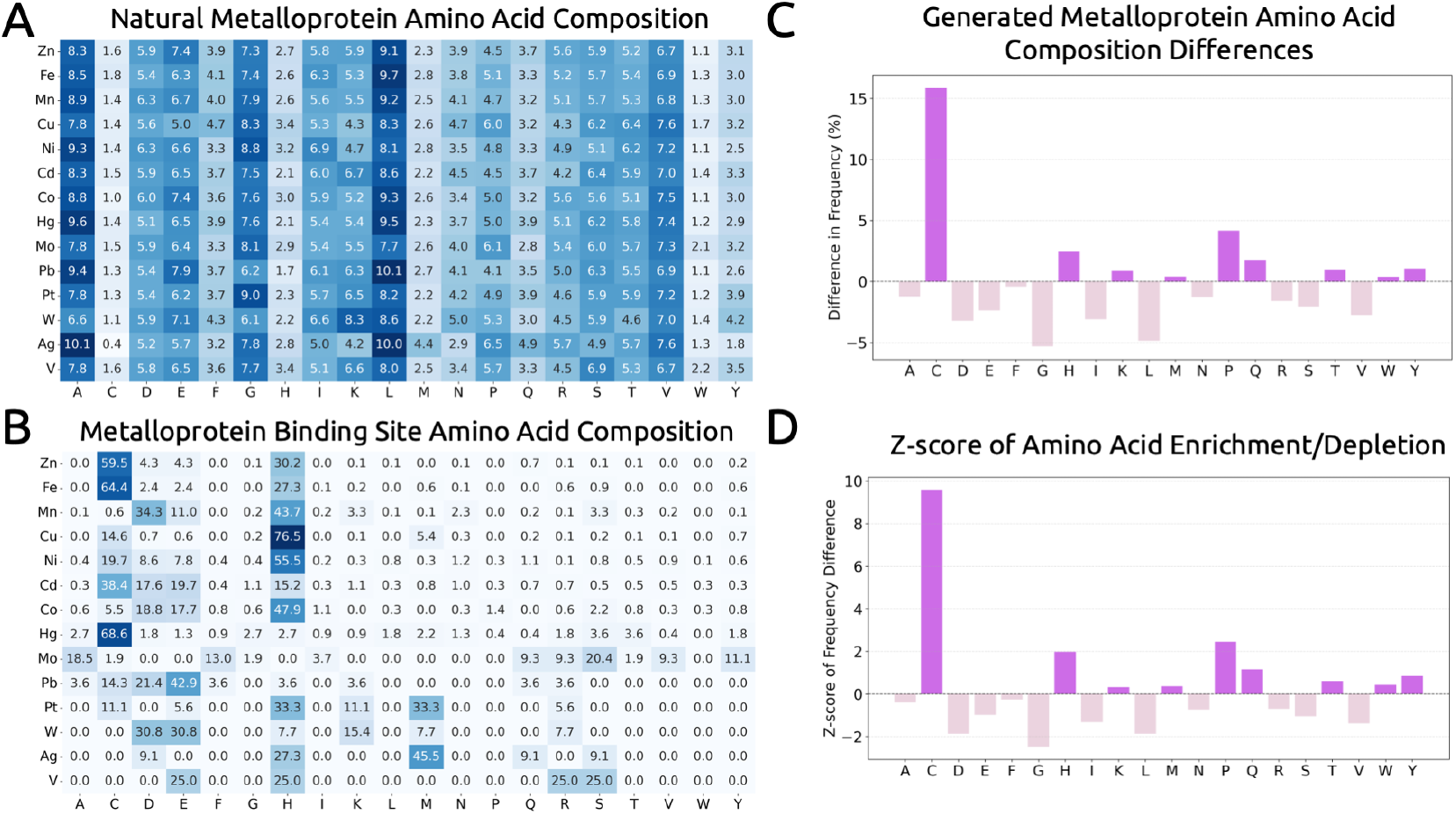
Amino Acid Composition Analysis between the MbPA database and the generated peptides. **A)** Aggregated natural amino acid composition from the MbPA database. **B)** Amino acid composition specifically at annotated metal-binding sites in MbPA (sites identified via PDB data). **C)** Composition differences (%) for generated *de novo* metalloproteins (Cd, Ni, Cu, and Zn) relative to MbPA. **D)** Corresponding Z-scores, computed by dividing each frequency difference by the standard deviation of that amino acid’s frequency in MbPA. The patterns broadly match those in (C).

**Figure S5:**
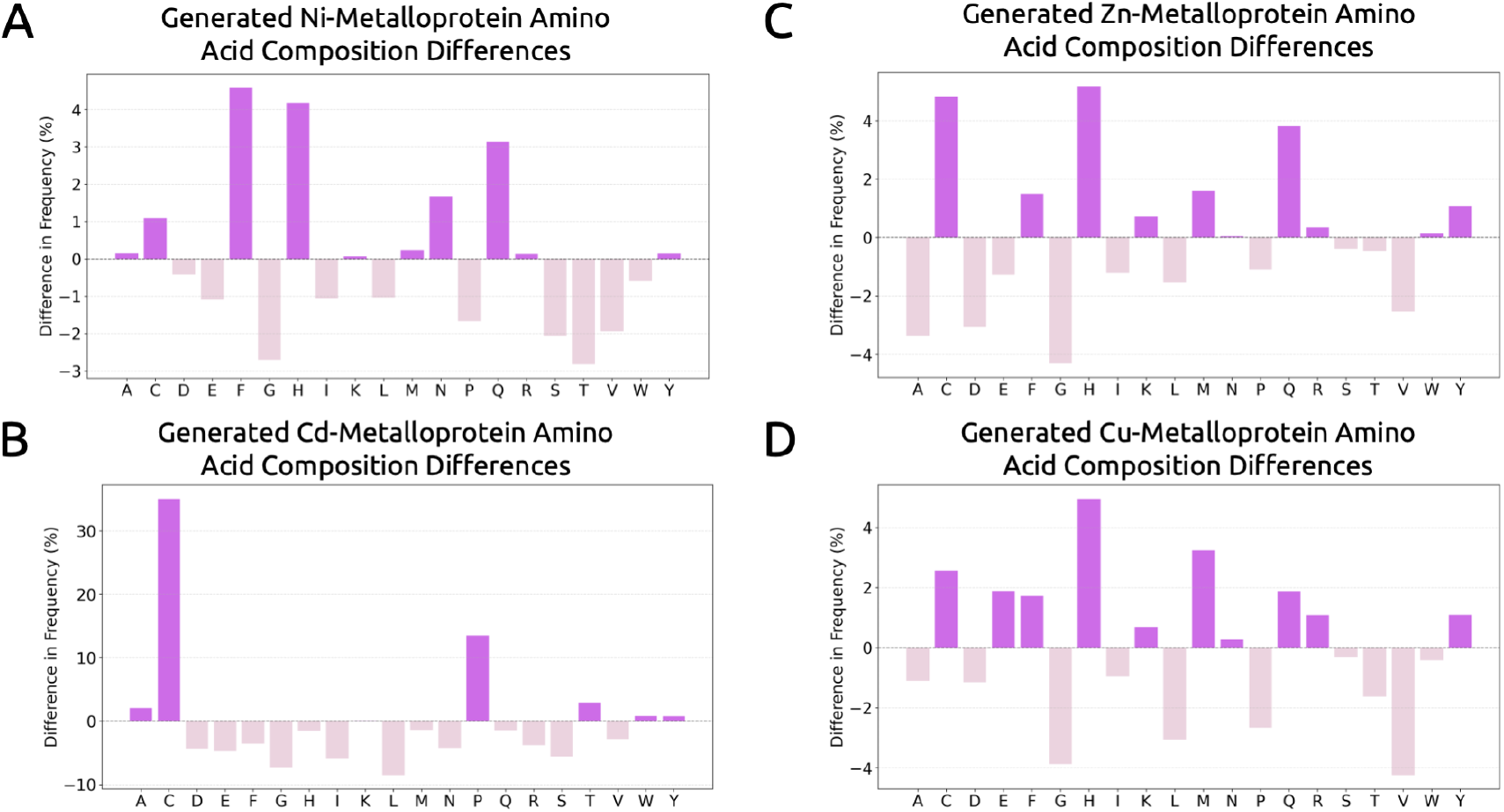
Metal-specific amino acid composition differences. **A)-D)** *De novo* composition differences (%) for Ni, Cd, Zn, and Cu, respectively, compared to the MbPA database (n=40, 890, 1000, 562).

**Table S3:**
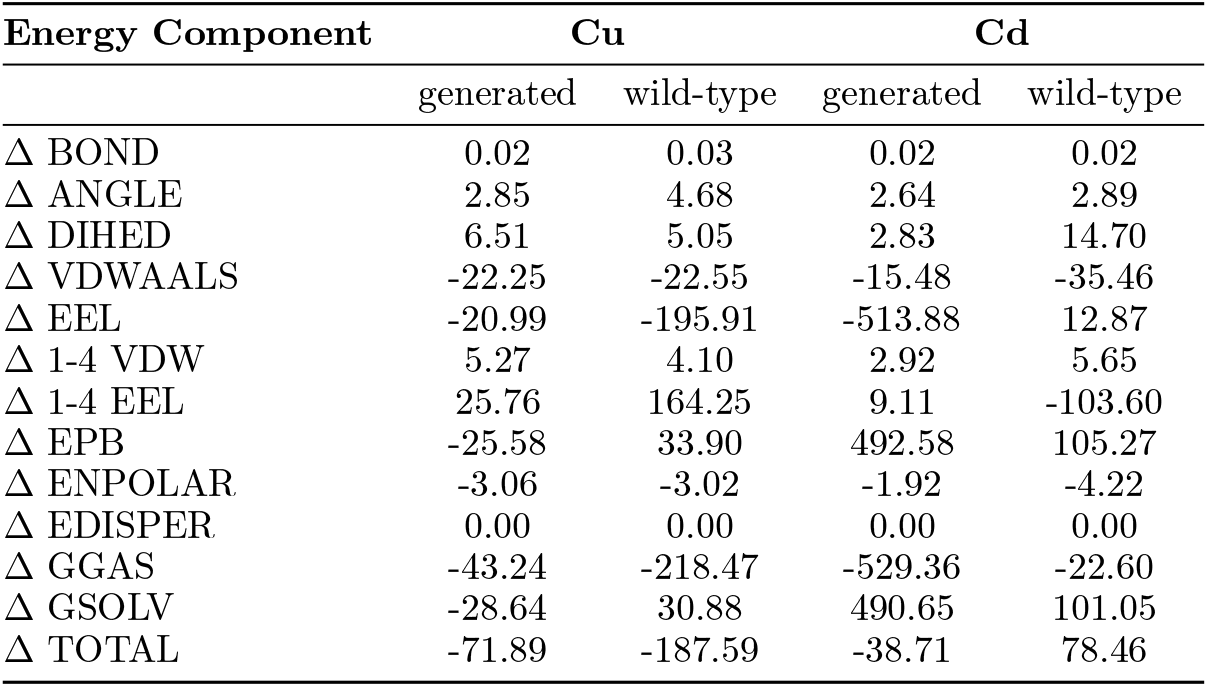
gmx_MMPBSA result. Delta (Complex - Receptor - Ligand). All units are reported in kcal/mol

## 4.2 Experimental Results

**Figure S6:**
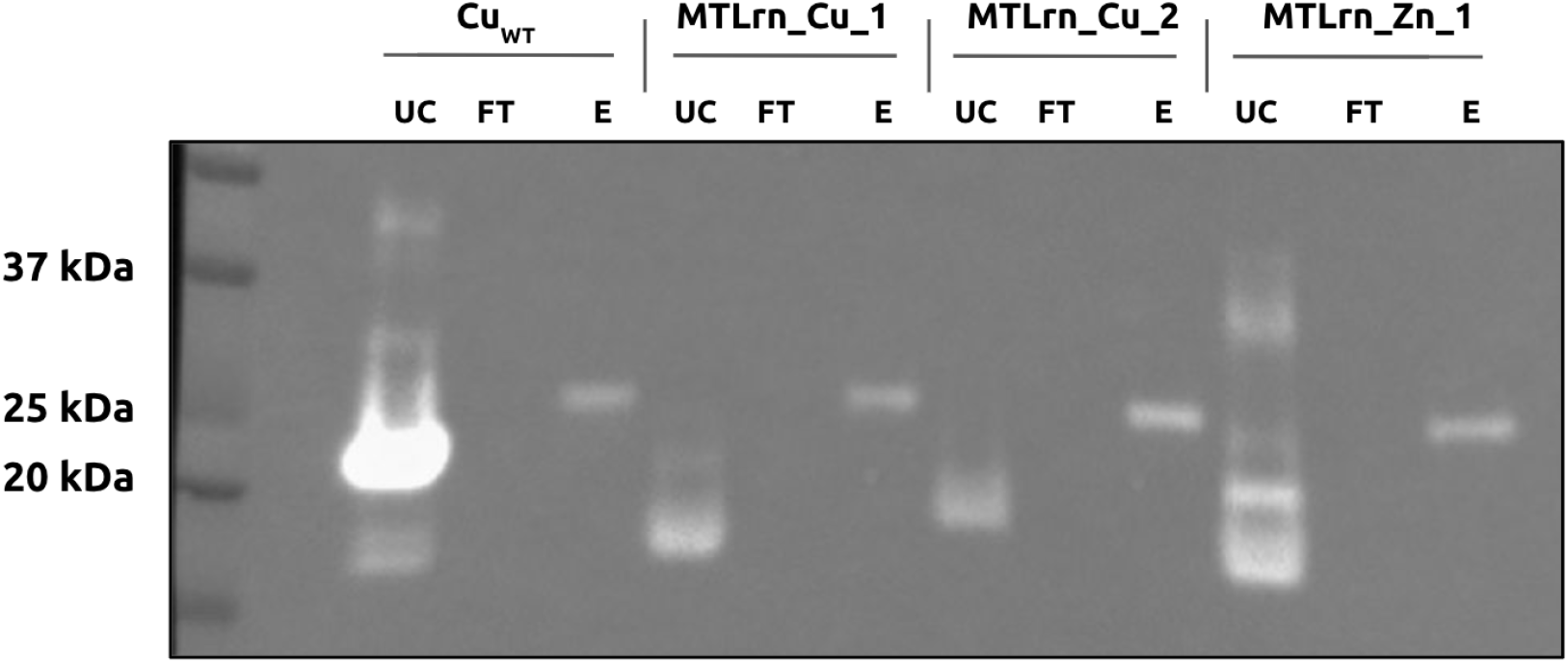
Anti-His Western blotting following reverse his-tag purification of cleaved constructs. Uncleaved (UC), flow-through (FT), and eluate (E) fractions of four SUMO-tagged constructs (Cu_WT_, MTLrn_Cu_1, MTLrn_Cu_2, MTLrn_Zn_1) were analyzed by anti-His Western blot to assess TEV cleavage efficiency and removal of the His-tag. A strong His signal is observed in UC lanes, confirming the presence of uncleaved His-tagged protein. After TEV protease treatment and reverse His-purification, cleaved proteins appear in the FT fractions with no His signal, indicating successful removal of the His-tag. A band in the E lanes corresponds to His-tagged TEV protease, which is retained on the Ni-NTA resin and eluted (∼28 kDa). Samples were loaded at 2 ug, transferred to PVDF membranes, and probed with HRP-conjugated anti-His antibody for chemiluminescent detection.

**Figure S7:**
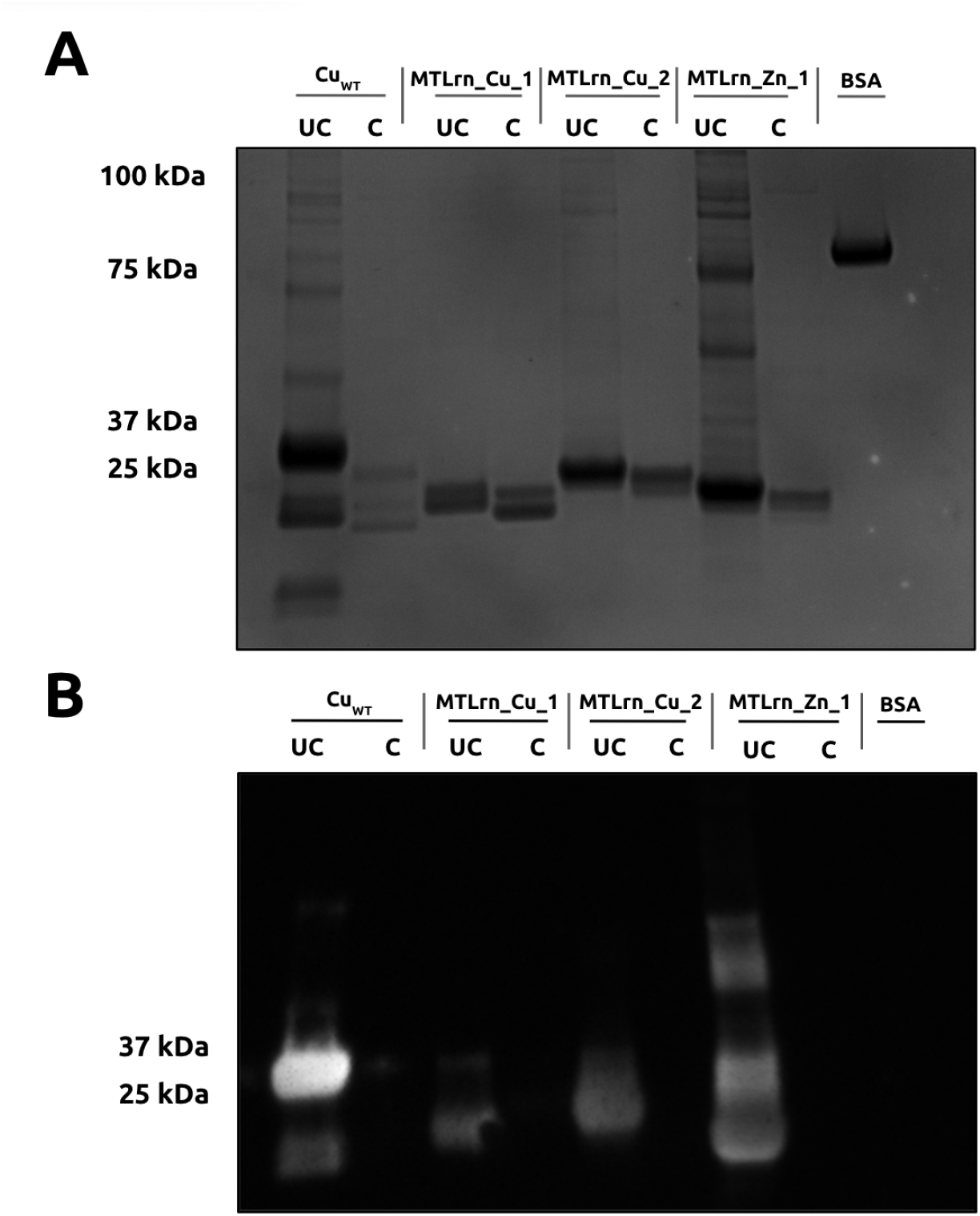
Confirmation of TEV-mediated cleavage via Coomassie-stained sodium dodecyl sulfate polyacrylamide gel electrophoresis (SDS-PAGE) and Anti-His Western Blotting. Constructs were analyzed before (UC, uncleaved) and after (C, cleaved) treatment with TEV protease. (A) SDS-PAGE showing a shift in molecular weight consistent with successful cleavage of the N-terminal 6xHis tag. (B) Anti-His Western blot performed on the same samples and gel loadings as in (A) confirms loss of the His-tag following TEV cleavage. His signal is present in UC lanes but absent in C lanes, indicating efficient cleavage. All samples (2 µg per lane) were denatured in LDS sample buffer with 5% *β*-mercaptoethanol, separated on 4–20% Tris-Glycine gels, and probed with HRP-conjugated anti-His antibody using chemiluminescent detection. ELISAs from Figure 4 were conducted on the above samples.

### 4.3 Extended Molecular Dynamics Analysis of *Cu*^2+^ and *Cd*^2+^ Binding Peptides

**Figure S8:**
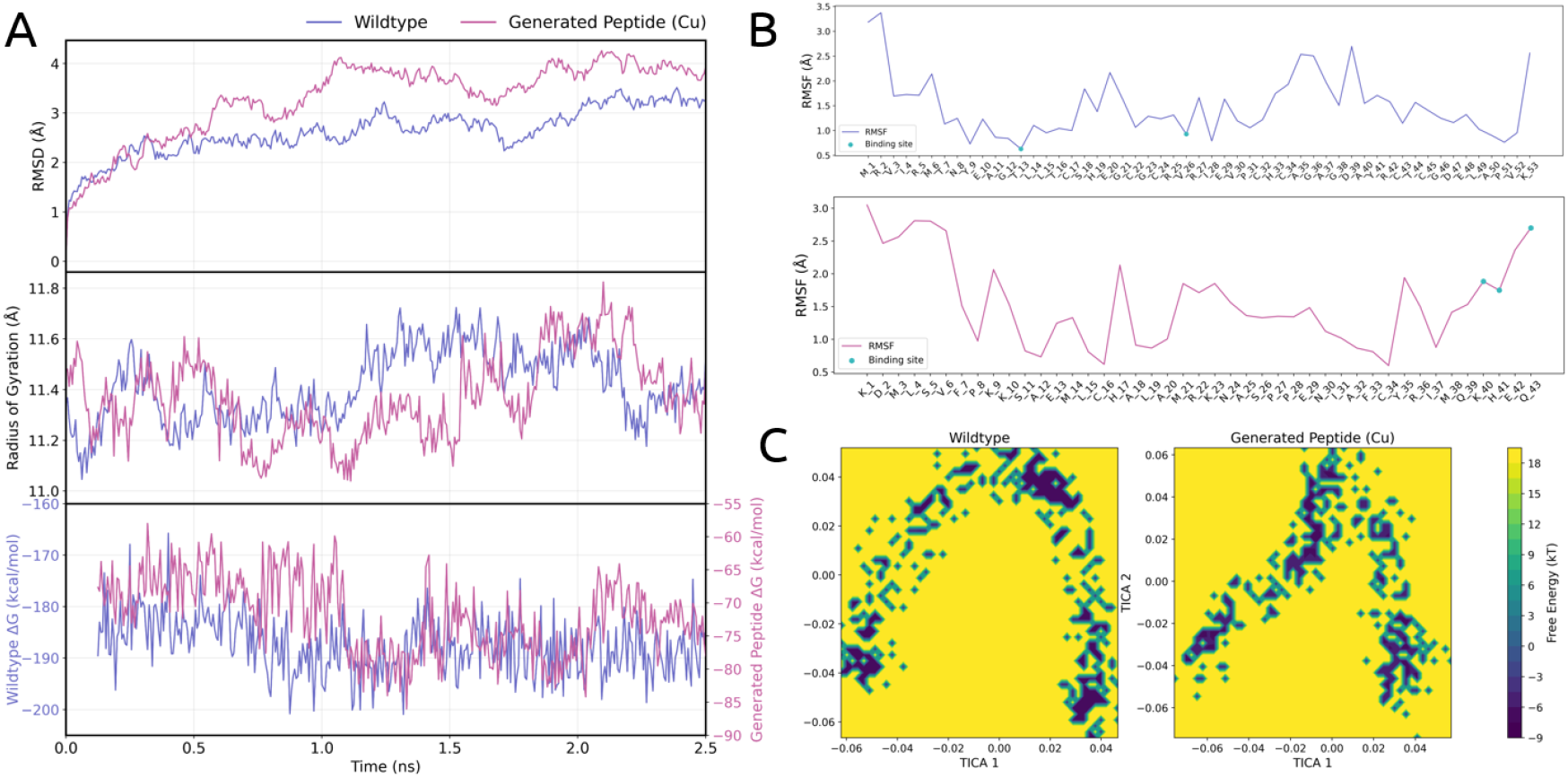
MM/PBSA analysis for selected Cu-binding peptide. **A)** Backbone RMSD, radius of gyration (Rg), and free energy surface for Metalorian-generated and wild-type Cu-binding peptides. The wild-type was selected to match the generated peptide in sequence length. Overall, structural stability and conformational preferences were comparable between the two systems. **B)** Pre-residue RMSF analysis. The generated peptide showed increased flexibility at the C-terminal region, where carboxyl-rich residues (e.g., K, H, Q) coordinate the metal. In contrast, the wild-type exhibited lower RMSF near its predicted binding site (T, V). Predicted coordinating residues, including canonical metal-binding residues such as C and H, generally exhibited low RMSF, indicating stable coordination. Elevated flexibility was observed only when binding sites overlapped with sequence termini, reflecting inherent mobility despite metal coordination. **C)** TICA landscapes of wild-type and Metalorian-generated peptides. Both systems exhibit a characteristic boomerang-shaped energy basin with minimal intermediate states, indicating similar conformational dynamics.

**Figure S9:**
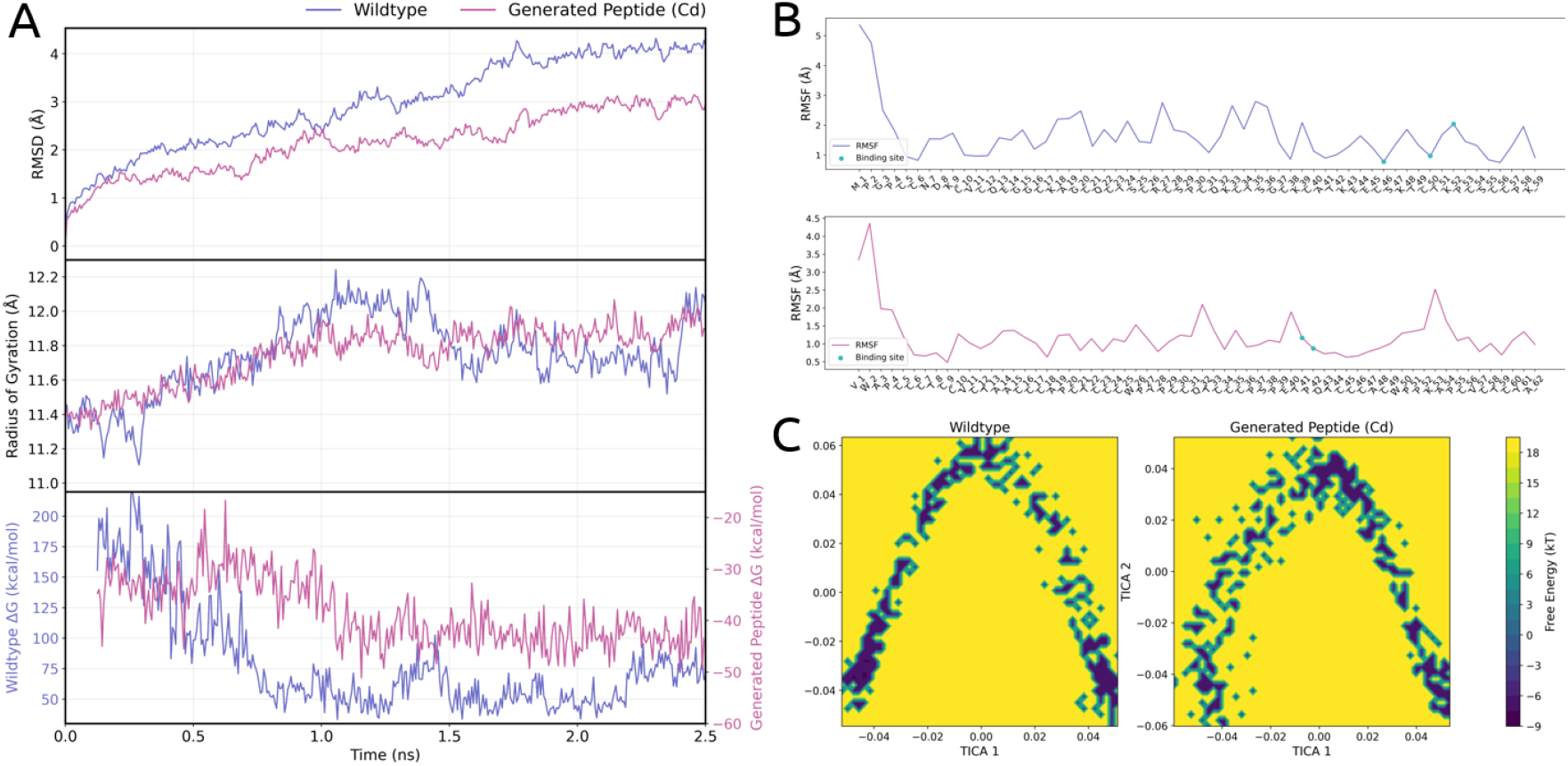
MM/PBSA analysis for selected Cd-binding peptide. **A)** Backbone RMSD, radius of gyration (Rg), and free energy surface for Metalorian-generated and wild-type Cd-binding peptides. The wild-type was selected to match the generated peptide in sequence length. Overall, structural features and dynamics were comparable between the two systems. The generated peptide exhibited a more favorable binding free energy (ΔTOTAL) compared to the wild-type, suggesting stronger metal coordination *in silico*. **B)** Per-residue RMSF analysis. Predicted binding sites—including canonical coordinating residues such as C and H—exhibited low RMSF, indicating stable coordination. **C)** TICA landscapes of wild-type and Metalorian-generated peptides. Both systems exhibit a characteristic boomerang-shaped energy basin with minimal intermediate states, suggesting similar dynamical pathways.

## 5 Pseudocode

### 5.1 EMA Thresholds Update

#### Algorithm 1 Classification Thresholds Update using EMA

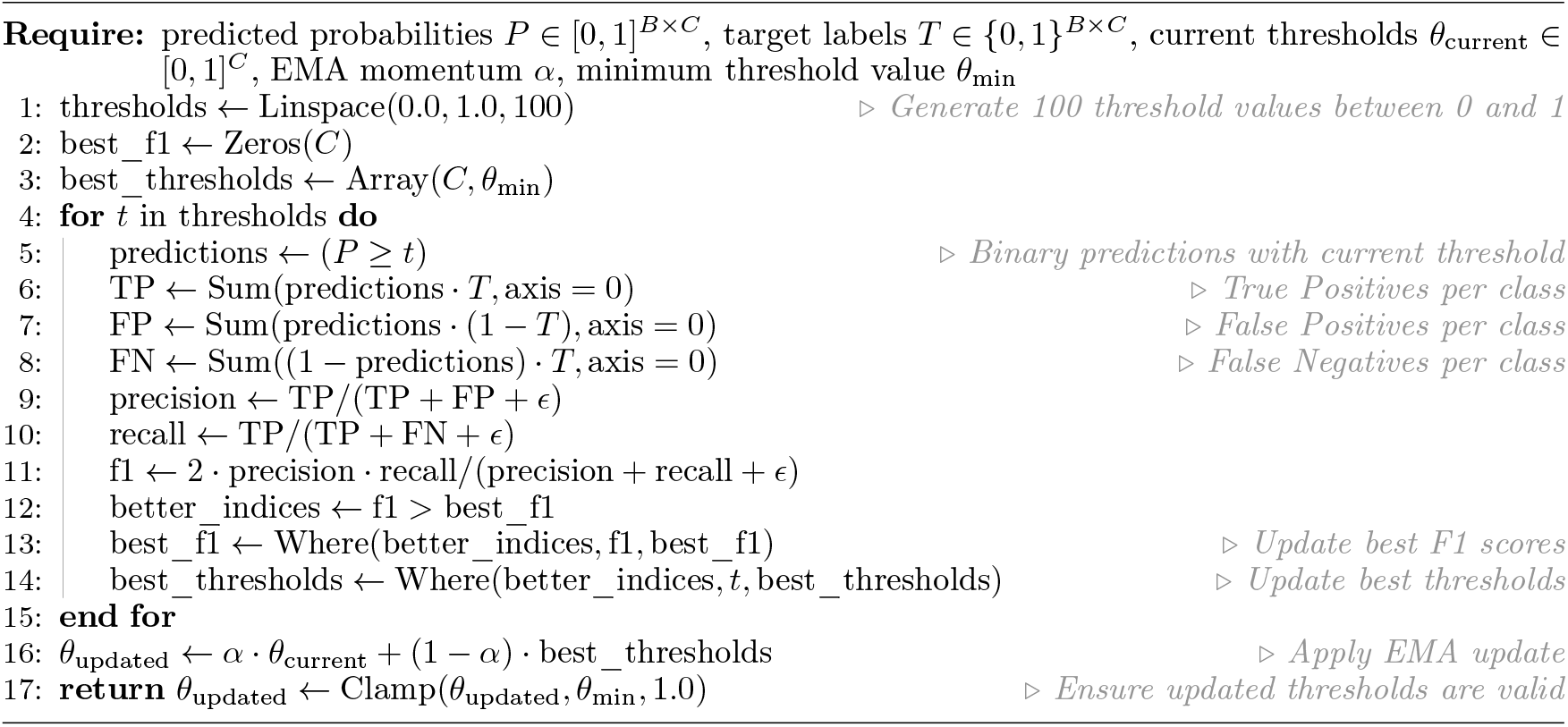

### 5.2 Adaptive Margin Triplet Loss

#### Algorithm 2 Adaptive Margin Triplet Loss

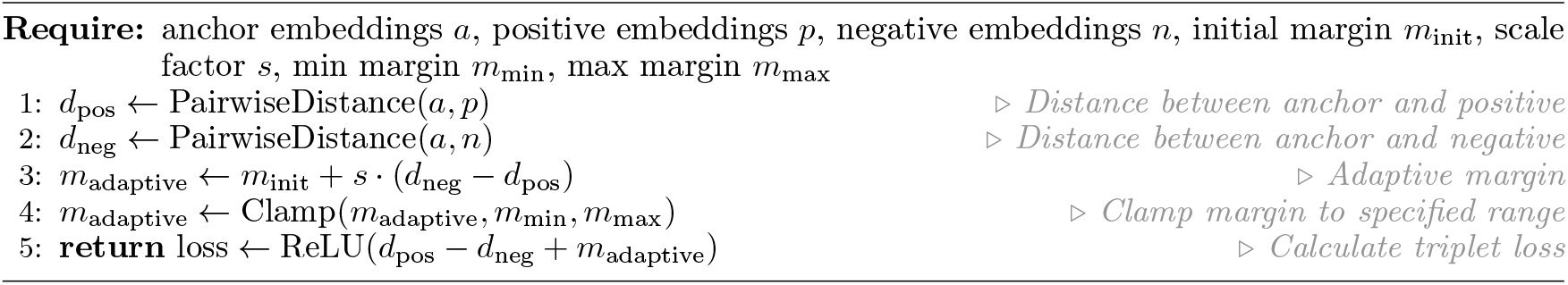

### 5.3 Pre-batching

#### Algorithm 3 Sequence Pre-Batching with Token Limit

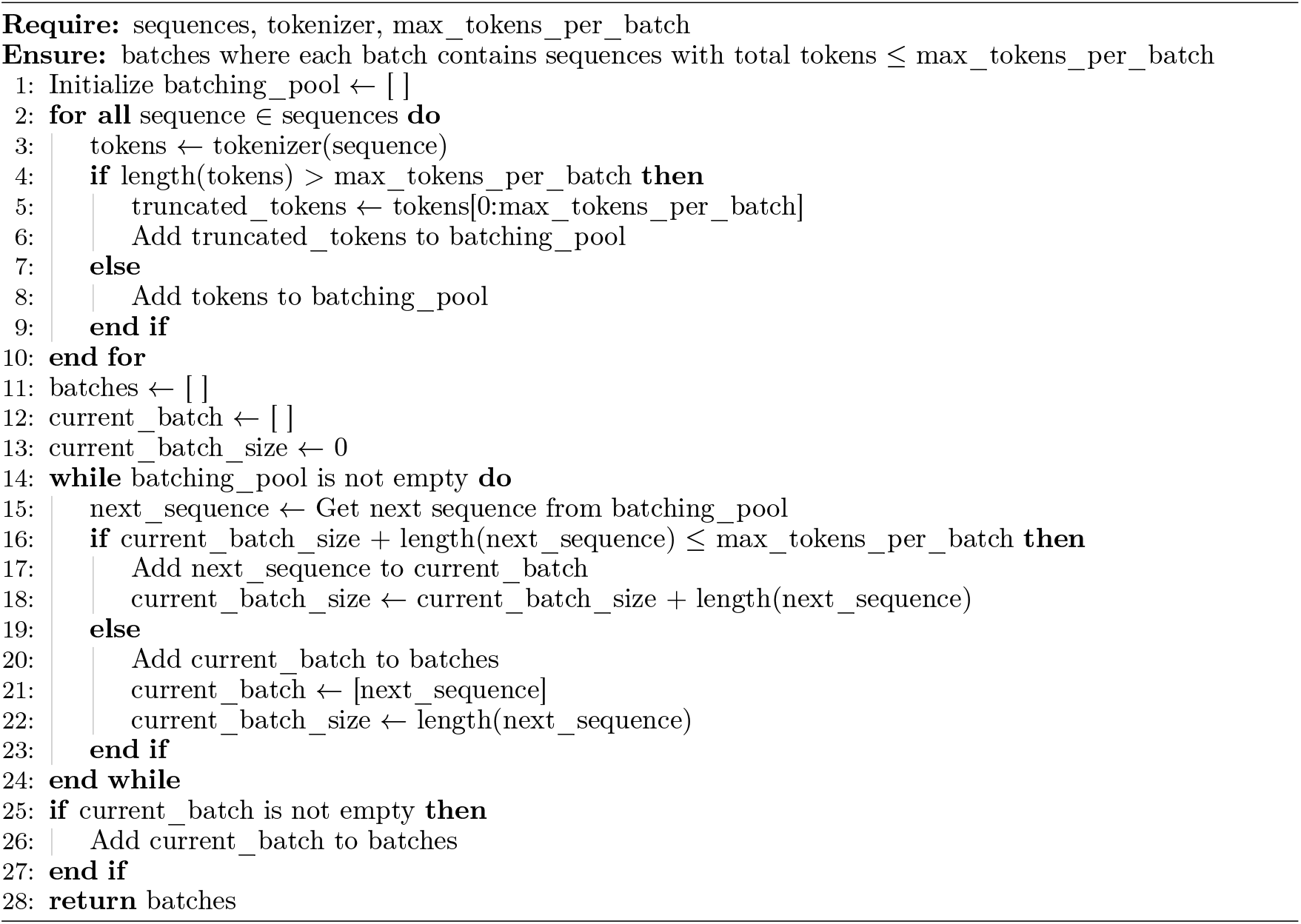

### 5.4 Negative training sample shifting

#### Algorithm 4 Progressive Shifting from Semi-Hard to Hard Negatives

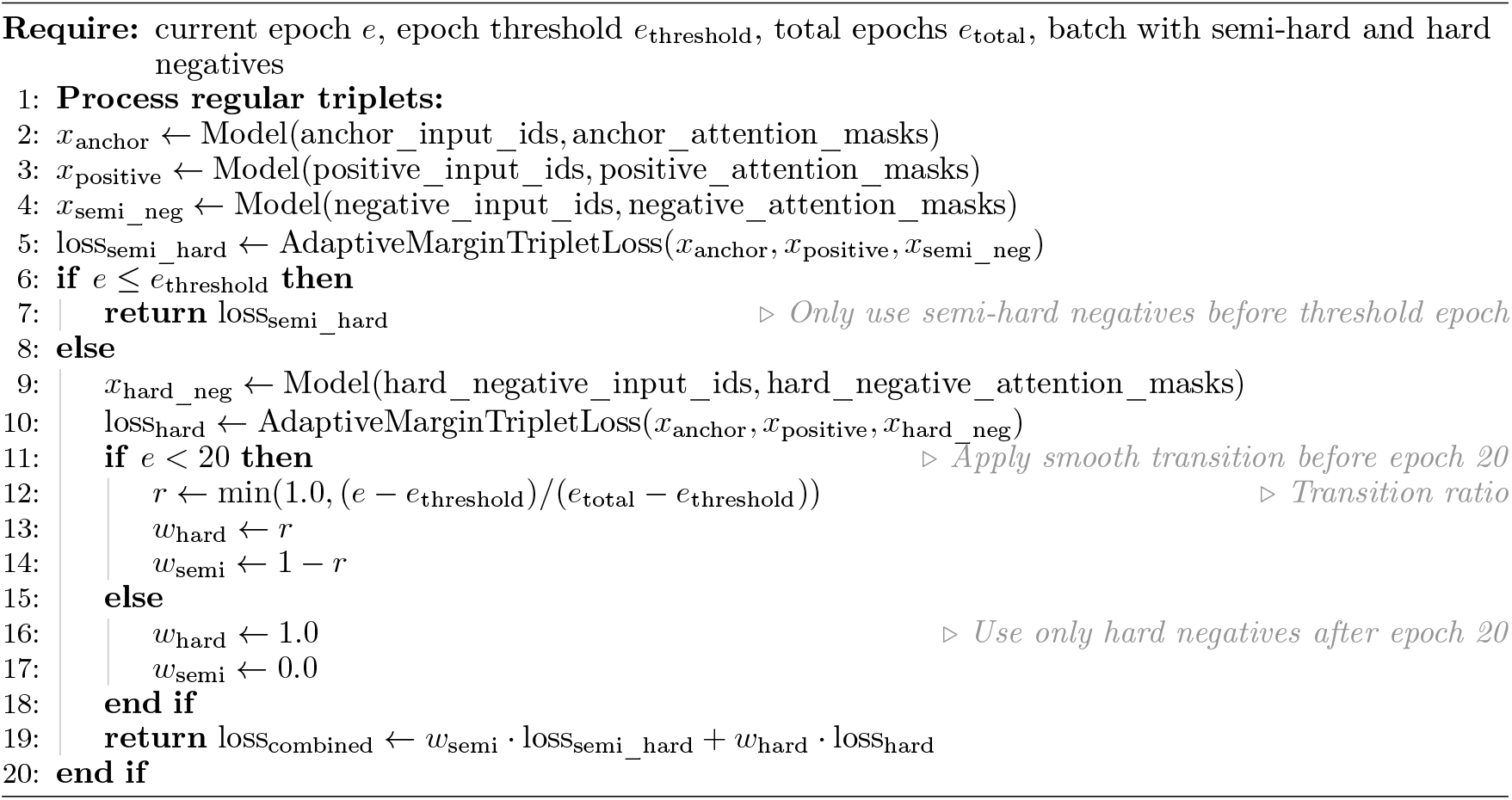

### 5.5 Sampling Pseudocode

#### Algorithm 5 Progressive Verification Sampling

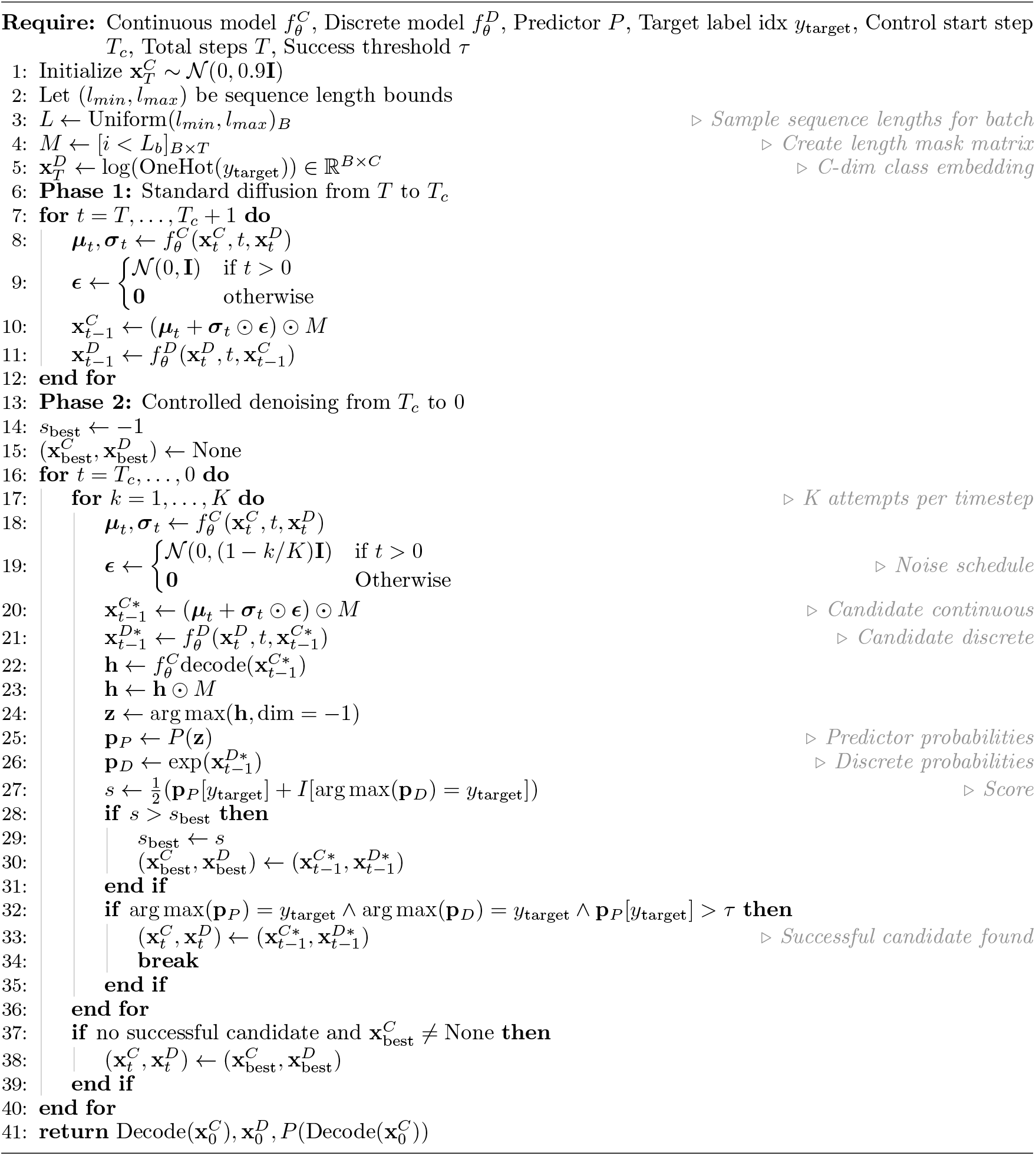

#### Algorithm 6 Gradient-Guided Label Sampling

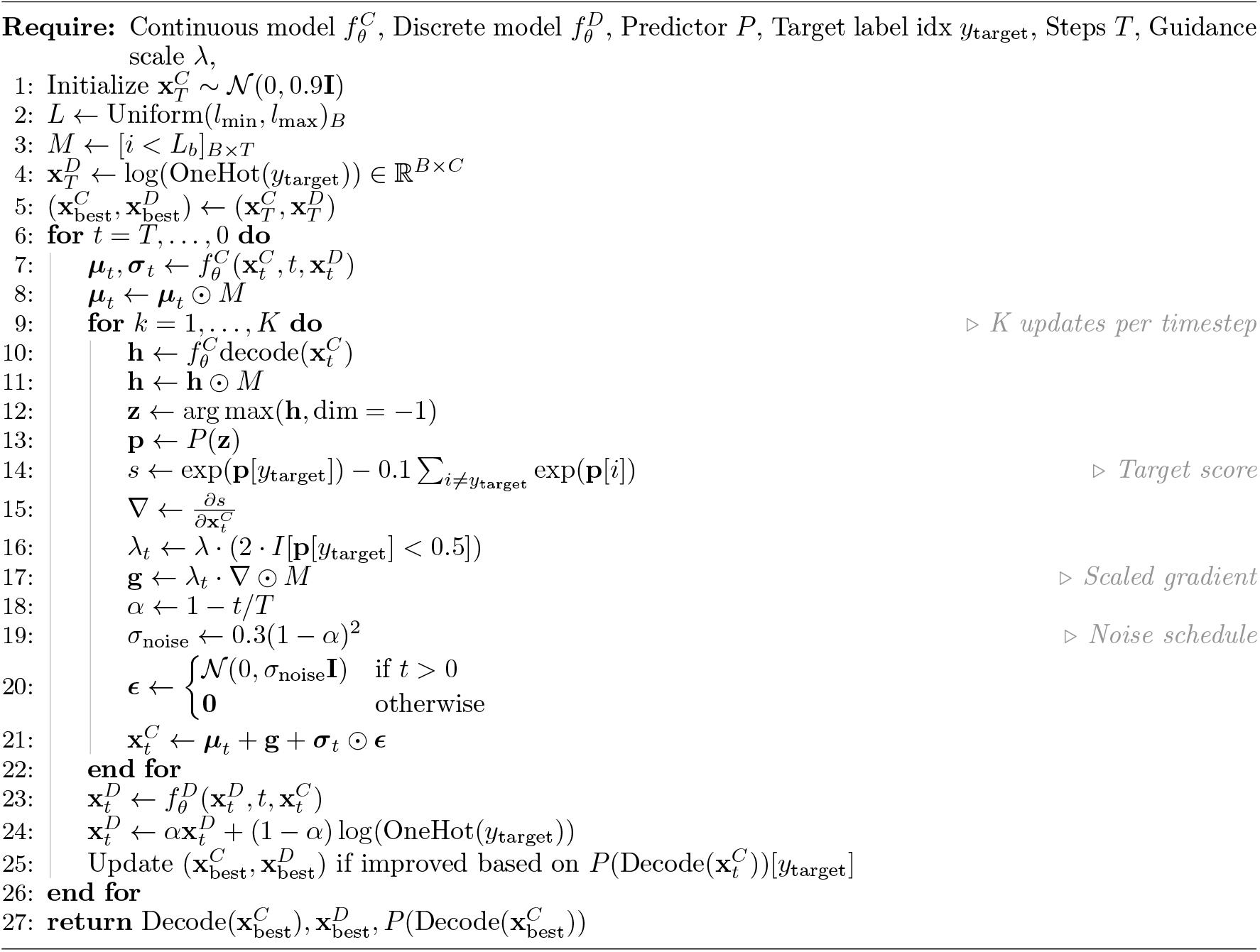

